# Senescent cells develop PDK4-dependent hypercatabolism and form an acidic microenvironment to drive cancer resistance

**DOI:** 10.1101/2022.08.29.505761

**Authors:** Xuefeng Dou, Qilai Long, Shuning Liu, Yejun Zou, Da Fu, Xue Chen, Qixia Xu, Changxu Wang, Xiaohui Ren, Guilong Zhang, Qiang Fu, Judith Campisi, Yuzheng Zhao, Yu Sun

## Abstract

Cellular senescence is a state of stable growth arrest, usually accompanied by development of the senescence-associated secretory phenotype (SASP). Although senescent cells remain metabolically active, little is known about their metabolic landscape and *in vivo* pathophysiological implications. Here we show that expression of the pyruvate dehydrogenase (PDH) inhibitory enzyme, pyruvate dehydrogenase kinase 4 (PDK4), is significantly upregulated in human senescent stromal cells. Preferentially expressed upon genotoxicity-induced senescence (GIS), PDK4 is negatively correlated with posttreatment survival of cancer patients. Upon cellular senescence, PDK4 shifts glucose metabolic flux from oxidative phosphorylation to aerobic glycolysis, causing enhanced lactate production and forming an acidic microenvironment. However, distinct from the cancer cell-featured Warburg effect, senescent cells maintain an intensive use of pyruvate through the tricarboxylic acid cycle (TCA), displaying increased respiration and redox activity, indicative of a special form of metabolic reprogramming. Conditioned media from PDK4^+^ stromal cells change global expression and promote malignancy of recipient cancer cells *in vitro* and accelerate tumor progression *in vivo*. Pharmacologically targeting PDK4 restrains the adverse effects of PDK4 in cell-based assays, while promoting tumor regression and extending posttreatment survival in preclinical trials. Together, our study substantiates the hypercatabolic nature of senescent cells, and reveals a metabolic link between senescence-associated acidic microenvironment and age-related pathologies including but not limited to cancer.

## Introduction

Cellular senescence was initially identified as a program characterized with loss of proliferative capacity after exhaustive passaging in culture, which is known as replicative senescence (RS) ^1^. Later studies demonstrated that senescence is indeed inducible by multiple types of inherent or environmental stresses, including oncogenic activation (oncogene-induced senescence, OIS) or therapeutic insults (therapy-induced senescence, TIS) ^2^. Senescent cells exhibit phenotypic alterations, such as morphological flattening, nuclear expansion, epigenetic reorganization and metabolic alterations ^3, 4^. They also display cell non-autonomous activities, particularly the active and continued secretion of numerous pro-inflammatory cytokines and chemokines, a phenotype termed the SASP ^5^. The SASP plays a context-dependent role in organismal aging and diverse age-related disorders ^4^. The net effect of the SASP is mostly detrimental in advanced life stages, as it does contribute to pathological incidence and clinical mortality. For example, the SASP promotes several hallmarks of cancer, while the elimination of senescent cells delays tumorigenesis ^6^. The number of senescent cells tends to increase with age in almost all mammalian species, resulting in significantly shortened healthspan and compromised lifespan ^7, 8^.

Biological experiments involving human cells and tissues, transgenic rodent models and pharmacological interventions consistently implicate senescent cells in age-related pathologies, while most of the detrimental effects of senescent cells can be attributed to the SASP, which is enriched with pro-inflammatory molecules ^9^. Single-cell profiling at both transcriptomic and proteomic levels suggest that senescent cells undergo intense metabolic reprogramming in order to maintain their cycle-arrested but viable status, and upregulate the expression of proteins essential to sustain the highly complex, dynamic and heterogeneous SASP ^3, 10^. In fact, several forms of metabolic stresses can both drive senescence and trigger the SASP. Many drivers of mitochondrial dysfunction contribute to cellular senescence, through disruption of cytosolic nicotinamide adenine dinucleotide (NAD^+^ and NADH), production of reactive oxygen species (ROS) and potentially other mechanisms. Of note, the mitochondrial dysfunction-associated senescence (MiDAS) phenotype lacks some pro-inflammatory components of the SASP, but instead exhibits a distinct set of SASP factors ^11^. MiDAS is primarily driven by the accumulation of cytosolic NADH, which is normally oxidized by mitochondria to NAD^+^. Decrease of the NAD^+^/NADH ratio inhibits the key glycolytic enzyme GAPDH, causing ATP depletion, AMPK activation and cell cycle arrest ^11^.

Senescent cells upregulate the protein level of the nicotinamide (NAM) salvage enzyme, namely nicotinamide phosphoribosyltransferase (NAMPT), while expression of the SASP markedly depends on NAD^+^ availability ^12^. Treating senescent cells with nicotinamide mononucleotide (NMN) can enhance the SASP, causing increased chronic inflammation and promoting development of inflammation-driven cancers ^13^. Thus, administration with NAD^+^-boosting supplements, such as nicotinamide riboside (NR) and NMN, may come at the cost of systemic side effects, such as enhancing chronic inflammation and cancer incidence. As a substrate to a group of diverse enzymes, including poly(ADP-ribose) polymerases (PARPs) and sirtuins, NAD^+^ is a critical nucleotide involved in oxidation-reduction reactions ^14^. However, NAD^+^ levels decrease during organismal aging and upon progeroid, causing metabolic dysfunction and a decline in physical fitness ^15, 16^. Recent studies further suggested that senescence-induced inflammation promotes CD38 accumulation in immune cells such as M1-like macrophages, while CD38 reduces levels of NMN and NAD^+^ through its ecto-enzymatic activity within metabolic tissues during chronological aging ^17, 18^. Despite these advances, a wide landscape of metabolic activities especially those correlated with glucose consumption and energy production, aspects essential to support the distinct protein synthesis machinery in senescent cells, remains hitherto lacking. In this study, we aimed to characterize the senescence-associated metabolism and uncovered that that pyruvate dehydrogenase kinase isoform 4 (PDK4), a pyruvate dehydrogenase (PDH) inhibitory kinase playing a key role in the control of metabolic flexibility of glucose, is significantly upregulated in senescent cells. Although displaying a reduced NAD^+^/NADH ratio, senescent cells produce an increased amount of pyruvate and lactate, metabolites correlated with elevated glycolysis. There is an inherent relationship between cellular senescence, PDK4 upregulation, acidic microenvironment and age-related diseases, specifically cancer. We propose a senescence-specific metabolic axis involving PDK4, a molecule that functionally supports metabolic reprogramming and may be exploited therapeutically to counteract aging and age-related pathologies.

## Results

### Genotoxic stress induces cellular senescence and PDK4 expression

Pyruvate enters the TCA cycle through PDH, while PDK molecules (PDK1-4) inhibit PDH activity and promotes the switch from mitochondrial oxidation to cytoplasmic glycolysis. PDK4 is located in the mitochondrial matrix and inhibits the PDH complex by phosphorylating its E1α subunit, thereby contributing to glucose metabolism regulation ^19^. Although predominantly expressed in the muscle and affects the metabolic fate of glucose during exercise, PDK4 is intensively studied in multiple cancer types ^20–22^. However, insights into PDK4 expression in normal tissue microenvironments and its inducibility in response to stressful insults remain limited. We recently noticed that the stromal cell line PSC27, which is of human prostate origin and comprising mainly fibroblasts but with a minor percentage of non-fibroblast cell lineages such as endothelial cells and smooth muscle cells, produces a large array of SASP factors upon exposure to cytotoxic insults specifically genotoxic chemotherapy or ionizing radiation ^23, 24^. Interestingly, PDK4 emerged as one of the upregulated factors, together with a list of typical SASP components, as previously revealed by our microarray profiling (Fig. 1a and Supplementary Fig. 1a) ^23^. To confirm the finding, we expanded by employing several alternative approaches to induce senescence, including replicative exhaustion (RS), overexpression of p16^INK4a^ (p16) and HRas^G12V^ (RAS), respectively. These treatment caused comprehensive cellular senescence, with an efficacy resembling that of DNA-damaging agents such as radiation (RAD), bleomycin (BLEO) and hydrogen peroxide (HP) (SA-β-Gal positivity and BrdU incorporation, Supplementary Fig. 1b-d). In each case, we observed a significant PDK4 induction (Fig. 1b-c).

**Fig. 1.**
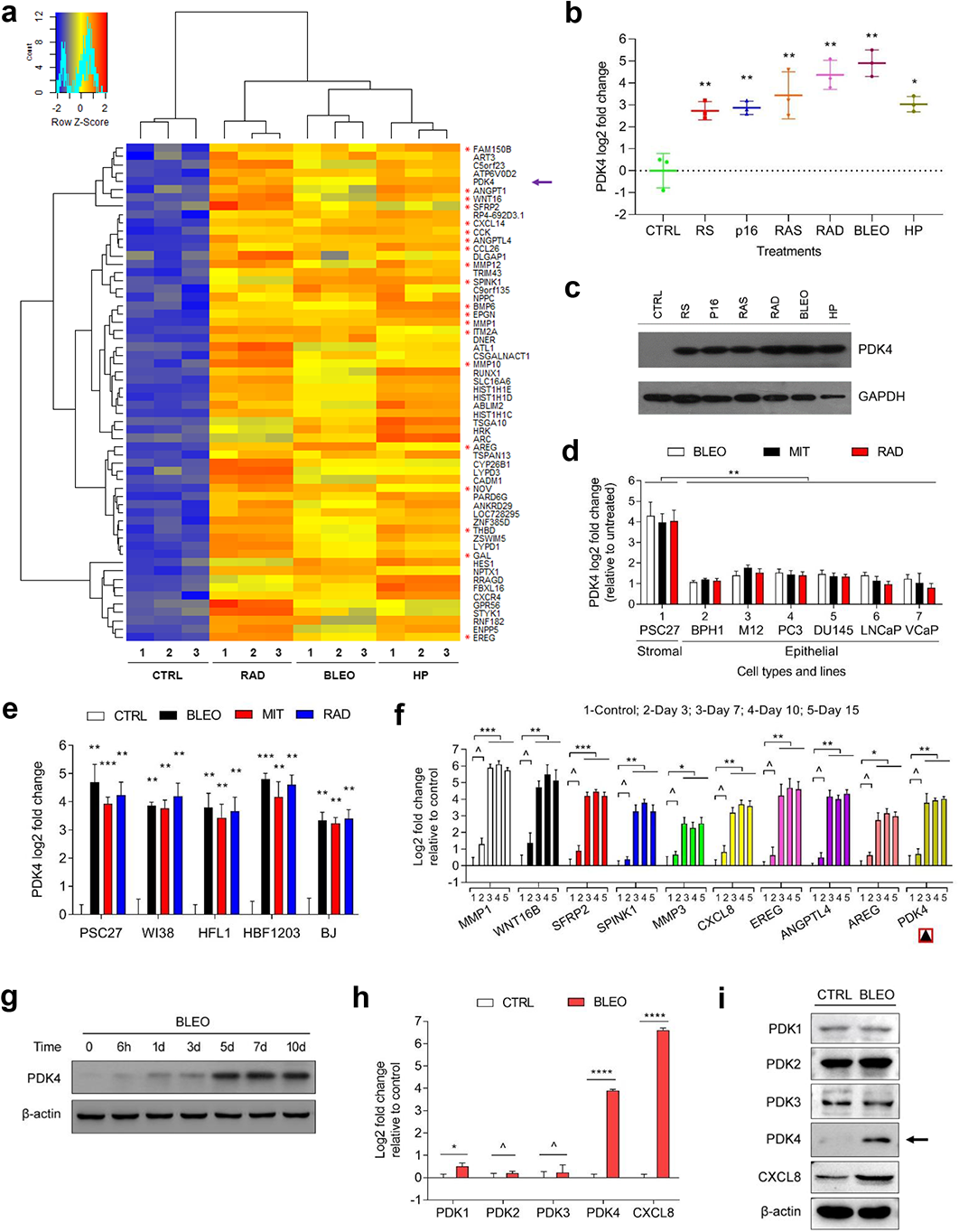
Genotoxicity induces senescence of human stromal cells which display upregulation of PDK4 and a full spectrum SASP. **a.** Transcriptome-wide profiling of gene expression changes in primary normal human prostate stromal cell line PSC27 by microarray. Cell lysates were collected for analysis 7 d after treatment. CTRL, control. RAD, radiation. BLEO, bleomycin. HP, hydrogen peroxide. Red highlighted, typical soluble factors of the SASP. Microarray data adapted from Sun et al. with permission from *Nature Medicine*, copyright 2012, Springer Nature ^23^. **b.** Quantitative RT-PCR to detect PDK4 expression after PSC27 cells were subject to individual treatment as indicated. Cell lysates were collected for measurement 7 d after establishment of stable cell sublines or completion of *in vitro* treatment. Signals normalized to CTRL. RS, replicative senescence. p16, lentiviral transduction of human tumor suppressor p16^INK4a^. RAS, lentiviral transduction of human oncogene HRAS^G12V^. **c.** Immunoblot analysis of PDK4 expression in stromal cells as delineated in (**b**). GAPDH, loading control. **d.** Comparative RT-PCR assay of PDK4 expression after treatment of PSC27 or prostate epithelial cells by agents as indicated. Cell lysates were collected for measurement 7 d after treatment. Signals normalized to CTRL. BPH1, M12, PC3, DU145, LNCaP and VCaP, human epithelial lines of prostate origin. **e.** Comparative RT-PCR assay of PDK4 expression in human stromal cells 7 d after treatments performed as indicated. WI38, HFL1, HBF1203 and BJ, human stromal lines of different origins. **f.** A time course RT-PCR assessment of the expression of PDK4 and a set of typical SASP factors (MMP1, WNT16B, SFRP2, SPINK1, MMP3, CXCL8, EREG, ANGPTL4 and AREG) after drug treatment of PSC27 cells *in vitro*. Numeric numbers indicate the individual days after treatment (indexed at the top line). **g.** Immunoblot measurement of PDK4 expression at protein level at the individual timepoints as indicated. β-actin, loading control. **h.** Comparative appraisal of human PDK family expression at transcript level in PSC27 cells after BLEO treatment. Signals normalized to untreated sample per gene. CXCL8, experimental control as a hallmark SASP factor. **i.** Immunoblot assessment of the expression of PDK4 family members at protein level after BLEO treatment. β-actin, loading control. Data are representative of 3 independent experiments. *P* values were calculated by Student’s *t*-test (**b**, **e**, **f**, **h**) and one-way ANOVA (**d**). ^, *P* > 0.05. *, *P* < 0.05. **, *P* < 0.01. ***, *P* < 0.001. ****, *P* < 0.0001.

Expression analysis of several cell lines of human prostate origin suggested that stromal cells are indeed more PDK4-inducible than cancer epithelial cells, implying a special mechanism that supports PDK4 production in prostate stromal cells posttreatment (Fig. 1d). Data from several additional fibroblast lines consistently supported a robust induction of PDK4 upon genotoxic treatment by anticancer agents (Fig. 1e). Notably, the transcript expression pattern of PDK4 largely phenocopied that of a group of hallmark SASP factors including MMP1, WNT16B, SFRP2, SPINK1, MMP3, CXCL8, EREG, ANGPTL4 and AREG, which exhibited a gradual increment until entering a platform within 7-8 days after treatment (Fig. 1f-g). Among the human PDK family (PDK1-4 isozymes), PDK4 appeared to be the only member that is readily inducible by genotoxic stress, with a tendency similar to that of CXCL8, an index of the SASP expression (Fig. 1h-i).

### PDK4 expression in stroma predicts adverse clinical outcomes post-chemotherapy

Experimental data derived from *in vitro* assays prompted us to further determine whether PDK4 induction occurs within the tumor microenvironment (TME), a pathological entity where a plenty of benign stromal cells reside. We first chose to analyze clinical samples of a cohort of prostate cancer (PCa) patients who developed primary tumors in the prostate and underwent chemotherapeutic regimen involving genotoxic agents such as mitoxantrone (MIT). Surprisingly, PDK4 was found markedly expressed in prostate tissues of these patients after chemotherapy, but not before (Fig. 2a). Basically in line with our *in vitro* data, upregulated PDK4 was generally localized in the stroma, in a sharp contrast to the adjacent cancer epithelium which had limited or no staining.

**Fig. 2.**
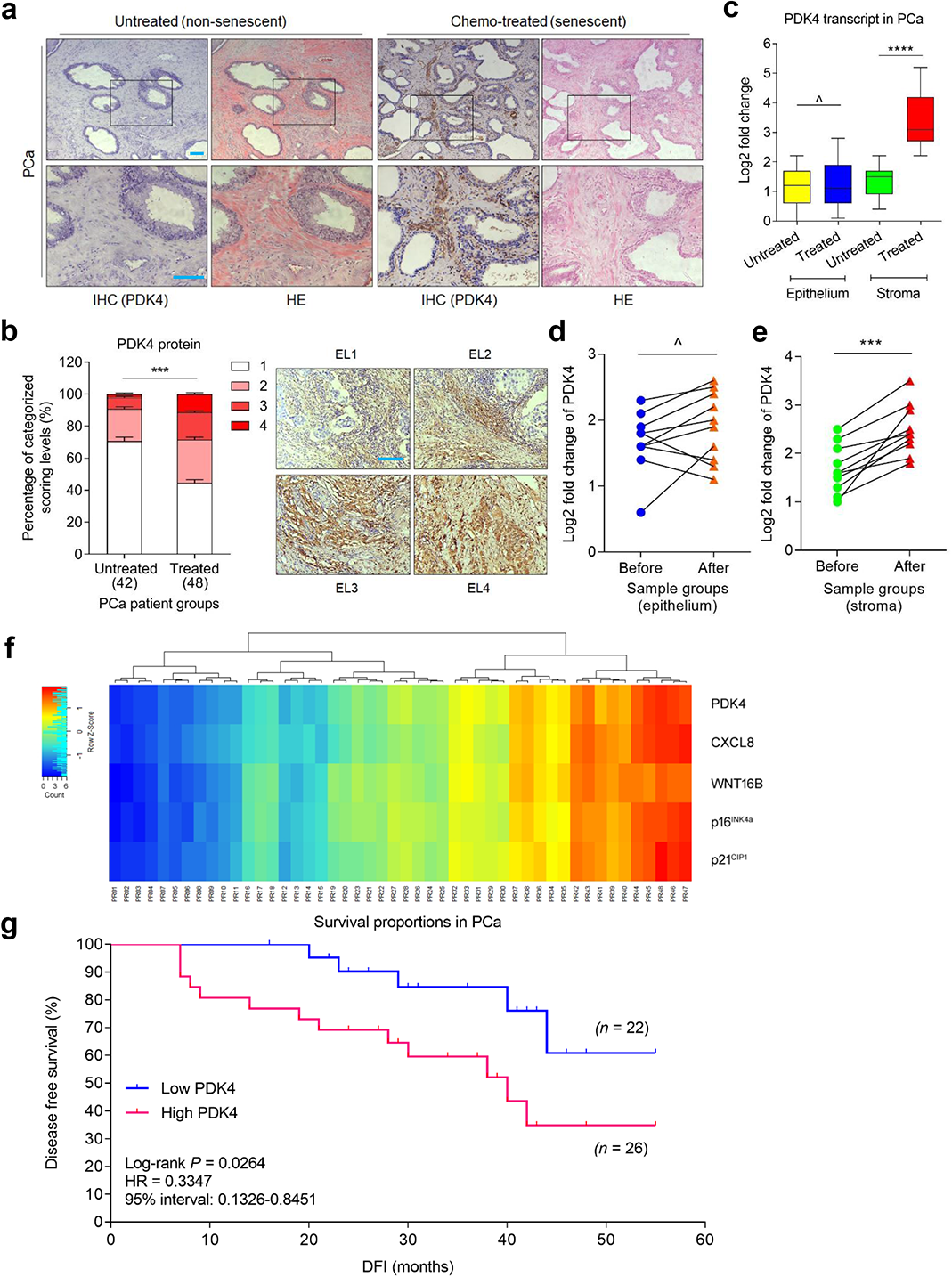
PDK4 is expressed in human prostate stroma after chemotherapy and negatively correlated with posttreatment survival. **a.** Representative images of PDK4 expression in biospecimens of human prostate cancer patients after histological examination. Left, immunohistochemical (IHC) staining. Right, hematoxylin and eosin (HE) staining. Left, untreated; right, chemo-treated. Rectangular regions selected in the upper images *per* staining are amplified into the lower images. Scale bars, 100 μm. **b.** Pathological assessment of stromal PDK4 expression in PCa samples (untreated, 42 patients; treated, 48 patients). Patients were pathologically assigned into 4 categories *per* IHC staining intensity of PDK4 in the stroma. 1, negative; 2, weak; 3, moderate; 4, strong expression. Left, statistical comparison. Right, representative images of each category regarding PDK4 signals. EL, expression level. Scale bar, 100 µm. **c.** Boxplot summary of PDK4 transcript expression by qRT-PCR analysis upon laser capture microdissection (LCM) of cells from tumor and stroma, respectively. Signals normalized to the lowest value in the untreated epithelium group, with comparison performed between untreated (42 patients) and treated (48) samples *per* cell lineage. For cells of either epithelium or stroma origin, samples from 10 patients out of untreated and treated groups were randomly selected for further analysis and parallel comparison. **d.** Comparative analysis of PDK4 expression at transcription level between epithelial cells collected before and after chemotherapy. Each dot represents an individual patient, with the data of “before” and “after” connected to allow direct assessment of PDK4 induction in the same individual patient. **e.** Comparative analysis of PDK4 expression at transcription level between stromal cells collected before and after chemotherapy. Presentation follows the manner described in (**d**). **f.** Pathological correlation between PDK4, CXCL8, WNT16B, p16^INK4a^ and p21^CIP1^ in the stroma of PCa patients after treatment. Scores were from the assessment of molecule-specific IHC staining, with expression levels colored to reflect low (blue) via modest (turquoise) and fair (yellow) to high (red) signal intensity. Columns represent individual patients, rows different factors. Totally 48 patients treated by chemotherapy were analyzed, with scores of each patient averaged from 3 independent pathological readings. **g.** Kaplan-Meier analysis of PCa patients. Disease free survival (DFS) stratified according to PDK4 expression (low, average score < 2, blue line, n = 22; high, average score ≥ 2, purple line, n = 26). DFS represents the length (months) of period calculated from the date of PCa diagnosis to the point of first time disease relapse. Survival curves generated according to the Kaplan–Meier method. HR, hazard ratio. Data in all bar plots are shown as mean ± SD and representative of 3 biological replicates. *P* values were calculated by Student’s *t*-test (**c**, **d**, **e**), two-way ANOVA (**b**) and log-rank (Mantel-Cox) test (**g**). ^, *P* > 0.05. *** *P* < 0.001. **** *P* < 0.0001.

PDK4 production in patient samples pre-*versus* post-chemotherapy was quantitatively measured by a pre-established pathological assessment procedure, which allowed precise evaluation of the expression of a target protein *per* immunohistochemistry (IHC) staining intensity (Fig. 2b). Transcript analysis upon laser capture microdissection (LCM) of cell lineages from primary tissues suggested that PDK4 was more readily induced in the stromal rather than cancer cell subpopulations (*P* < 0.0001 versus *P* > 0.05) (Fig. 2c). To substantiate PDK4 inducibility *in vivo*, we profiled a subset of PCa patients whose pre- and post-chemotherapy biospecimens were both accessible, and found remarkably upregulated PDK4 in the stroma, but not cancer epithelium, of each individual post-chemotherapy (Fig. 2d-e). We noticed the dynamics of PDK4 expression in the damaged TME largely in parallel to that of CXCL8 and WNT16B, two canonical SASP components (Fig. 2f). Expression pattern of these factors were largely consistent with that of senescence markers including p16^INK4a^ and p21^CIP1^ in tumor foci, suggesting an inherent correlation of PDK4 induction with cellular senescence and the SASP (Fig. 2f). Of note, Kaplan-Meier analysis of PCa patients stratified according to PDK4 expression in tumor stroma suggested a significant but negative correlation between PDK4 protein level and disease-free survival (DFS) in the treated cohort (*P* < 0.05, log-rank test) (Fig. 2g).

The distinct pathological properties of PDK4 in PCa were subsequently reproduced by an extended study, which was designed to recruit clinical cohorts of human breast cancer (BCa) patients (Supplementary Fig. 2a-d). Implicating the functional roles of PDK4, such as working as a critical regulator of epithelial-to-mesenchymal transition (EMT) and drug resistance of human cancers ^21^, data from gene expression profiling interactive analysis (GEPIA) with the cancer genome atlas (TCGA) and genotype-tissue expression (GTEx) databases indicated that PDK4 expression in cancer cells *per se* is associated with the poor prognosis of some, but not all cancer types (Supplementary Fig. 2e-f). Thereby, in contrast to former studies that mainly focused on PDK4 expression in cancer cells *per se*, our data consistently suggest that PDK4 induction in tumor stroma may act as an SASP-associated independent predictor of clinical prognosis, holding the potential to be exploited for stratifying the risk of disease relapse and clinical mortality of posttreatment patients. Given such a pathological relevance, it is reasonable to speculate that PDK4 production by the stroma may have a causal role in senescence-related conditions, such as cancer progression.

### Senescent cells exhibit a distinct profile of glucose metabolism

A typical feature of cancer cells is the ability of reprogramming energy metabolism to fuel their expansion and survival, while enhanced mitochondrial function plays important roles in tumor development ^25^. One of the major hallmarks of senescent cells is that they remain metabolically active and synthesize a plethora of protein factors (SASP) with a capacity to affect other cells of the host microenvironment locally or systemically ^26^. Former studies on the metabolism of cellular senescence demonstrated that levels of both glucose consumption and lactate production are elevated during senescence ^27^. While increased expression of glucose transporter and glycolytic enzymes during cellular senescence was observed, to date relevant data were mostly derived from cancers such as lymphomas and melanomas, or senescent cells induced by activation of oncogenes such as BRAF^V600E^ ^28, 29^. In contrast, the metabolic feature of glucose, a major energy source of senescent cells, and the influence of such a metabolic profile on surrounding tissue homeostasis, remain largely underexplored and merits in-depth understanding.

Glucose is the primary carbon source to the tricarboxylic acid (TCA) cycle, followed by glutamate and aspartate (non-protonatable amino acids as glutamine or asparagine, respectively) as secondary sources (Fig. 3a) ^30^. We first interrogated the metabolic pattern of glucose upon uptake by senescent cells, as glucose is supposed to act as a principal contributor to the TCA cycle when cells enter senescence, a stage that is considered metabolically active ^31^. Experimental data from assessment of mitochondrial dynamics and cellular bioenergetics with the XF24 Extracellular Flux analyzer (Seahorse bioanalyzer) indicated significantly elevated glycolytic activity in senescent human stromal cells, as reflected by enhanced production of metabolites including but not limited to dihydroxyacetone phosphate (DHAP), glyceraldehyde-3-phosphate (GAP) and 3-phosphoglycerate (3-PG) (Fig. 3b). Increased levels of GAP and 3-PG imply further utilization of a number of middle metabolites, such as citrate, α-ketoglutarate, glutamate, succinate, fumarate and malate in TCA cycle, which was indeed substantiated by data from metabolic profiling with the Seahorse bioanalyzer (Fig. 3c and Supplementary Fig. 3a-f). Thus, bioactivities of both glycolysis and TCA cycle were significantly enhanced in senescent cells, as reflected by an overall profiling with assays of stable isotope labelling with a uniformly labelled U-^13^C_6_ glucose tracer and fractioning of metabolites derived from labelled glucose (Fig. 3d). We further noticed that these metabolic changes were accompanied by remarkable perturbations in mitochondrial ultrastructure of senescent cells, particularly the enlarged size and abnormal shape as revealed by transmission electron microscopy (TEM), a phenomenon indicative of ultrastructural damage of mitochondria and suggesting potential mitochondrial dysfunction associated with oxidative stress upon cellular senescence (Fig. 3e). These observations are largely consistent with former studies regarding abnormal mitochondrial phenotypes including their mass, dynamics and structure in senescent cells ^32^.

**Fig. 3.**
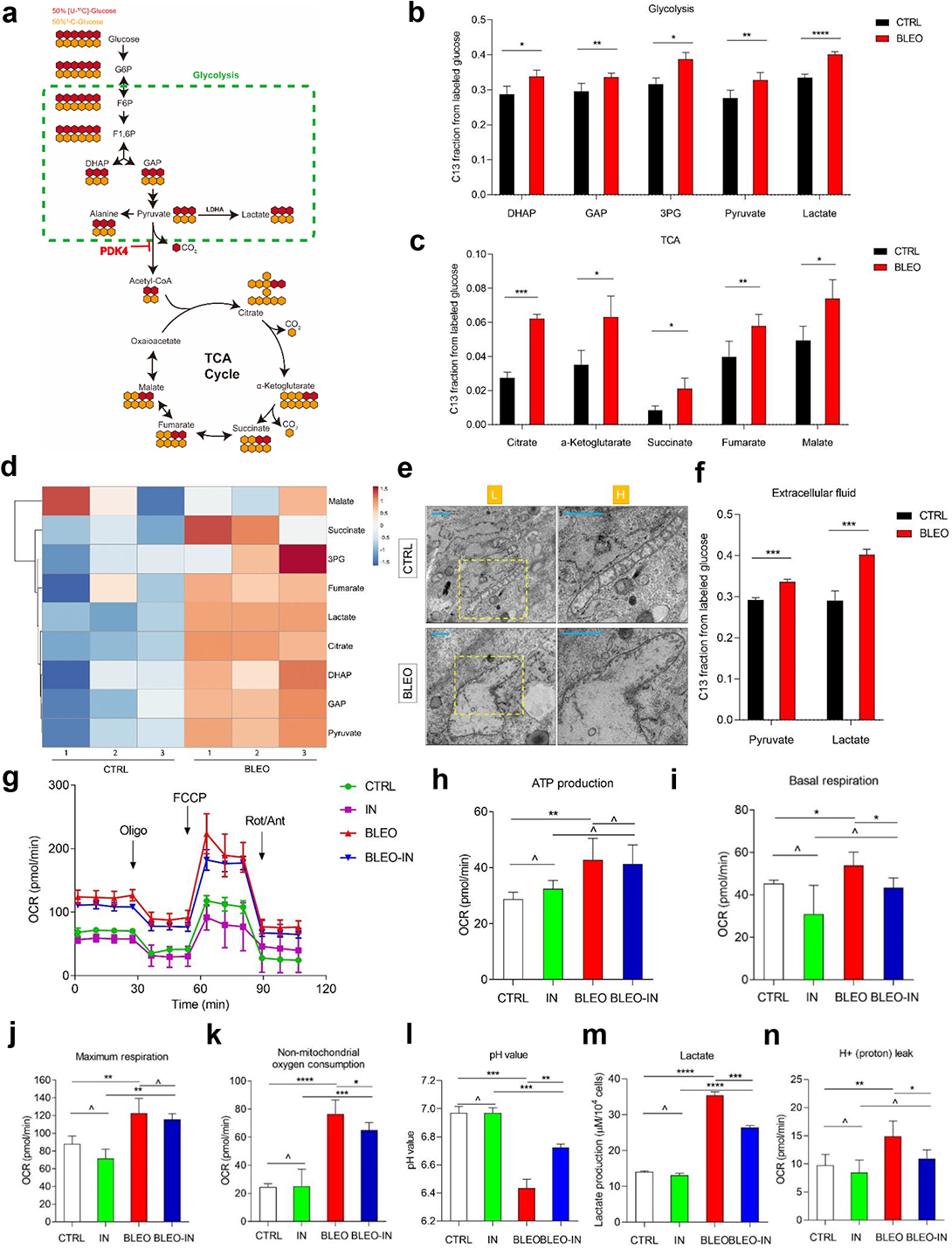
Senescent cells display a glucose metabolic profile distinct from their normal and cancer counterparts. **a.** A schematic molecular roadmap briefly outlining the landscape of glucose metabolism in mammalian cells. **b.** Partial metabolic profiling (glycolysis) of senescent cells induced by BLEO (TIS) and incubated with uniformly labeled [U-^13^C_6_]-glucose. Results from analysis of metabolites including DHAP, GAP, 3PG, pyruvate and lactate are shown. **c.** Partial metabolic profiling (TCA cycle) of senescent cells induced by BLEO (TIS) and incubated with uniformly labeled [U-^13^C_6_]-glucose. Results from analysis of metabolites including citrate, α-ketoglutarate, succinate, fumarate and malate, are presented. TCA, tricarboxylic acid cycle. **d.** Heatmap depicting the changes of glucose catabolism-associated metabolites as measured for senescent cells by the XF24 Extracellular Flux Analyzer. **e.** Representative TEM images showing the ultrastructural profile of mitochondria in human stromal cells. L, low resolution. H, high resolution. Scale bars, 1.0 μm. **f.** Measurement of extracellular fluids with XF24 extracellular flux analyzer. Pyruvate and lactate were assayed as indicated. **g.** OCR of stromal cells was measured using an XF24 extracellular flux analyzer. Briefly, 1.5μM oligomycin, 0.5 μM FCCP and 0.5 μM Rot/Ant were injected sequentially into each well. Glycolysis rate was calculated as (maximum rate measured before Oligo injection) minus (last rate measured before glucose injection). All Seahorse data were normalized with cell numbers, with metabolic parameters automatically calculated by WAVE software equipped in Seahorse. OCR, oxygen consumption rate. Oligo, oligomycin. FCCP, carbonyl cyanide 4-(trifluoromethoxy) phenylhydrazone. Rot, rotenone. Ant, antimycin. IN, PDK4-IN (PDK4 inhibitor) **h.** Measurement of ATP production by stromal cells. ATP production was measured as (last rate measurement before Oligomycin injection) minus (minimum rate measurement after Oligomycin injection). **i.** Assessment of basal respiration as an essential element of the senescence-associated metabolism program. **j.** Examination of maximal respiration as another fundamental element of the senescence-associated metabolism program. **k.** Assessment of non-mitochondrial oxygen consumption in stromal cells. **l.** Measurement of pH values in stromal cells. **m.** Determination of lactate production in stromal cells. **n.** Examination of the leak of H+ (proton) from mitochondria of stromal cells. Data in all bar plots are shown as mean ± SD and representative of 3 biological replicates. *P* values were calculated by Student’s *t*-test (**b, c, f, h-n**). ^, *P* > 0.05. * *P* < 0.05. ** *P* < 0.01. **** *P* < 0.0001.

We next measured the levels of extracellular fluids. Strikingly, the amounts of both pyruvate and lactate released to the extracellular space were considerably enhanced in senescent cells relative to their control counterparts (Fig. 3f). These changes were accompanied by alterations in oxygen consumption rate (OCR) and extracellular acidification rate (ECAR), suggesting elevated metabolic activities associated with glucose utilization (Fig. 3g and Supplementary Fig. 3g-i). Correspondingly, we observed elevated ATP production, basal respiration, maximum respiration in senescent cells, a tendency indicative of tight connection of TCA cycle and oxidative phosphorylation (OXPHOS) but further promoted when PDK4-IN-1, an anthraquinone derivative and a potent inhibitor of PDK4 (referred to as PDK-IN hereafter) ^33^, was applied to culture (Fig. 3h-j). However, treatment with PDK4-IN reversed the tendency of such changes in non-mitochondrial oxygen consumption, pH fluctuation, lactate production and H^+^ (proton) leak, with the overall metabolic data validated by principal component analysis (PCA) scores (PC1 *vs* PC2) (Fig. 3k-n and Supplementary Fig. 3j). There alterations occurred in parallel with expression changes of glucose uptake-associated molecules and metabolism-related enzymes including glucose transporter 1 (GLUT1), hexokinase 2 (HK2), lactate dehydrogenase A (LDHA), isocitrate dehydrogenase 2 (IDH2), isocitrate dehydrogenase 3 (IDH2), oxoglutarate dehydrogenase (OGDH) and citrate synthase (CS) (Supplementary Fig. 3k). Among them, HK2 and LDHA are glycolysis-related factors, while IDH2, IDH3, OGDH and CS are TCA cycle-associated enzymes. As overexpression of PDK4 *per se* caused neither cellular senescence nor the SASP (Supplementary Fig. 3l, m), we reasoned that the metabolic profile of senescent cells was correlated with and likely underpinned by expression of key factors involved in glucose consumption and linked with production of pyruvate, lactate and multiple other metabolites. Of note, elevated levels of glycolysis and oxidative phosphorylation were simultaneously observed, suggesting essentially reprogrammed glucose metabolism upon cellular senescence.

Former studies reported that in contrast to proliferating cells, senescent cells exhibit increased glucose transporter and glycolytic enzyme expression levels after chemotherapeutic treatment ^28^, a tendency largely confirmed by our experimental data (Supplementary Fig. 3k). Steady state glucose concentrations tend to be higher in senescent cells as compared to their control, suggesting an elevated glucose avidity upon senescence. These findings are essentially confirmed by our metabolomics profiling, which underscores the global catabolic nature of senescence-associated metabolic alterations. Taken together, senescent cells develop a distinctive hypermetabolic phenotype characterized of enhanced glycolysis, TCA activity and ATP-boosting oxidative phosphorylation. Increased energy production is a common denominator of senescent cells, which exhibit a specific utilization of energy-generating metabolic pathways, a phenomenon largely reminiscent of the ‘Warburg effect’ typically observed in cancer cells capable of performing a non-oxidative breakdown of glucose ^28^.

### Senescent cells shapes the formation of an acidic microenvironment *via* PDK4

Data from previous studies indicated that senescent cells are in a hypermetabolic status, more specifically, display a hypercatabolic nature ^28^, thus promoting us to interrogate whether these cells have a glucose uptake capacity distinct from that of proliferating cells. To address this, we performed another set of metabolic assays. Not surprisingly, a significant increase of glucose uptake by senescent cells was observed, although with the tendency preferentially detected upon TIS (Fig. 4a). Further, the pH of conditioned media (CM) from senescent cells was markedly decreased, a property that again seemed to be more dramatic for cellular senescence induced by genotoxic agents (Fig. 4b). Given the results indicative of an elevated acidification potential as revealed by the ECAR assay (Supplementary Fig. 3g), we reasonably speculated the extracellular formation of an acidic microenvironment by senescent cells, whose metabolism appeared to be distinctly reprogrammed and was characterized with increased secretion of acidic metabolites. Our data suggested that senescent cells can generate an increased amount of lactate, which was higher than their cycling control counterparts (Supplementary Fig. 4a). Multiple studies reported that cancer cells have increased lactate production, OCR level and ATP production, a series of metabolic changes correlated with enhanced glycolysis ^34, 35^. Strikingly, we found that many relevant activities of senescent cells were even higher than those of cancer cells selected as of the same organ origin (herein, prostate), such as PC3 and DU145, although with several key features manifesting changes evidently opposite to those of examined cancer lines (Supplementary Fig. 4a-f).

**Fig. 4.**
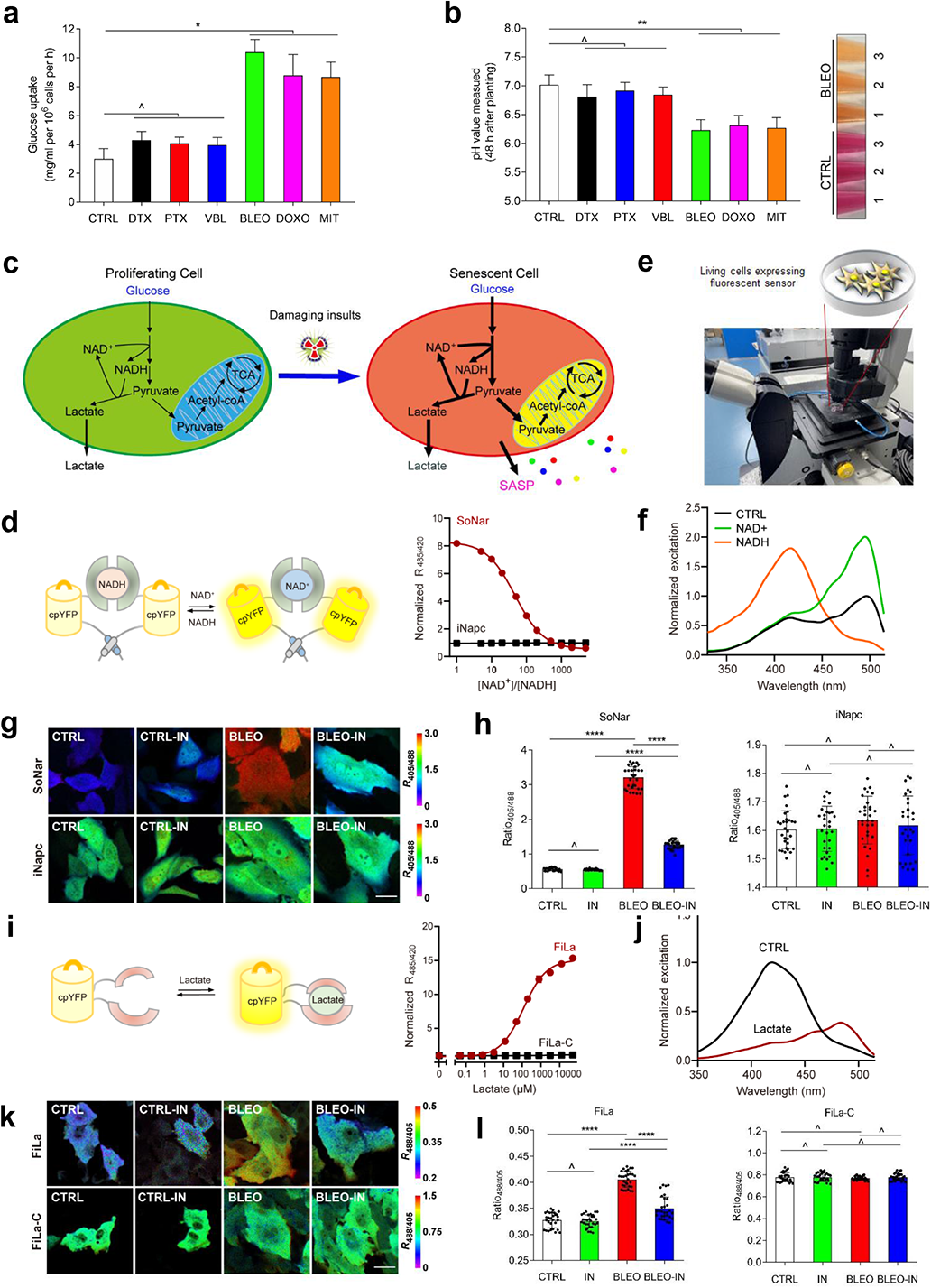
Senescent stromal cells display changes of NAD^+^/NADH metabolism and form an acidic microenvironment *via* PDK4-mediated production of lactate. **a.** Measurement of glucose uptake of PSC27 cells upon senescence induction by therapeutic agents as indicated. **b.** Examination of the pH value of human stromal cells treated as described in (**a**). Right inlet, representative images of conditioned media collected from proliferating and BLEO-induced senescent cells, respectively. **c.** Schematic illustration of potential changes in cell metabolic activities specifically fluctuations of the NAD^+^/NADH ratio during senescence induced by damaging insults. **d.** Graphic model for design of SoNar, which is a fusion of cpYFP and the NADH-binding domain of T-Rex. Binding of NAD^+^ or NADH induces changes in the protein conformation and fluorescence. Right, fluorescence ratios plotted against the NAD^+^/NADH ratio at 400 μM total nicotinamide adenine dinucleotide. Fluorescence ratios were normalized to the control condition in the absence of nucleotides (n = 3). **e.** Technical overview for *in vitro* imaging of living cells stably expressing the fluorescent sensor with confocal laser-scanning microscopy. **f.** Excitation spectra of purified SoNar in the control condition (black), and after addition of 20 μM NAD^+^ (green) or 20 μM NADH (orange), normalized to the peak intensity in the control condition. Emission was measured at 530 nm. **g.** Fluorescence imaging of SoNar in proliferating (CTRL) or bleomycin-induced senescent (BLEO) PSC27 cells, in the absence or presence of PDK inhibitor (IN) for 2 h, Scale bar, 20 μm. **h.** Quantification of SoNar or iNapc fluorescence in PSC27 cells (n = 30 cells). Left, SoNar. Right, iNapc. **i.** Schematic representation of molecular design for lactate sensor FiLa. The fluorescent protein cpYFP was inserted into a monomer of the lactate-binding bacterial protein LldR. Binding of lactate induces changes in protein conformation and fluorescence. Right, Lactate titration curves of FiLa and FiLa-C sensors. Data are normalized to the initial value (n = 3). **j.** Excitation spectra of purified FiLa in the control condition (black) and saturated with lactate (dark red). The excitation spectrum recorded at an emission wavelength of 530 nm has maxima at 425 and 490 nm. Data are normalized to peak intensity at 425 nm in the control condition. **k.** Fluorescence imaging of FiLa or FiLa-C in CTRL and BLEO-induced senescent PSC27 cells, in the absence or presence of PDK inhibitor (IN) for 2 h, Scale bar, 20 μm. **l.** Quantification of FiLa (left) and FiLa-C (right) fluorescence in PSC27 cells (n = 30 cells). Data in all bar plots are shown as mean ± SD and representative of 3 biological replicates. *P* values were calculated by Student’s *t*-test. ^, *P* > 0.05. * *P* < 0.05. ** *P* < 0.01. *** *P* < 0.001. **** *P* < 0.0001.

PDK4 is a key enzyme involved in the regulation of glucose and fatty acid metabolism as well as tissue homeostasis, while its overexpression inactivates the PDH complex by phosphorylating the targets and contributes to metabolic flexibility. We assessed the influence of PDK4 expression by transducing a PDK4 construct to human stromal cells, and noticed significantly altered metabolic profile including glucose uptake, lactate production and triglyceride (TG) production, although these changes were largely reversed upon genetic eliminated of PDK4 (Supplementary Fig. 4g-i). Of note, a decreased pH of the CM was observed in the case of PDK4 overexpression in proliferating cells, but subject to counteraction by PDK4 suppression (Supplementary Fig. 4j). We further measured these parameters with senescent cells induced by BLEO treatment, and found markedly increased glucose uptake, lactate production and TG production, but reduced pH of the CM (Supplementary Fig. 4k-n). However, almost all these metabolic changes were substantially reversed, when PDK4 was depleted from PSC27, except the TG levels, a case suggesting a PDK4-mediated antagonism against TG synthesis throughout the TCA cycle in senescent cells (Supplementary Fig. 4k-n). We noticed some factors functionally supporting glycolysis and TCA, including GLUT1, MCT4, HIF1α, PGK1, PGI, CS, IDH2, IDH3A and IDH3B were concurrently upregulated upon TIS, further indicating an overall enhancement of cell metabolism with the glucose as an energy source (Supplementary Fig. 4o).

NAD^+^ and its reduced form, NADH, are pivotal coenzymes for redox reactions and play critical roles in energy metabolism ^36^. The intracellular level of NAD^+^ is frequently altered during aging and upon age-related pathologies. We previously generated SoNar, an intensely fluorescent, rapidly responsive, pH-resistant and genetically encoded sensor for tracking subtle changes in cytosolic NAD^+^ and NADH redox states by imaging and quantifying the NAD^+^/NADH ratio in living cells and *in vivo* ^37^, but the metabolic profile NAD^+^ and NADH in senescent cells remains largely undefined (Fig. 4c). We first measured the intracellular NAD^+^/NADH redox state of PSC27 cells utilizing SoNar’s fluorescence (Fig. 4d-f). The data indicated a remarkably lower NAD^+^/NADH ratio upon TIS (evidenced by increased NADH/NAD^+^), a tendency that was essentially subject to reversal by the PDK4 inhibitor, suggesting a remarkably elevated reduction of NAD^+^ to NADH, a process accompanied by enhanced glycolysis (Fig. 4g, h). As another technical advancement, we designed FiLa, a highly responsive, ratiometric and genetically encoded lactate sensor to monitor the production and consumption of lactate at subcellular resolution (Fig. 4i-j). We observed a remarkable increase in cytosolic lactate upon cellular senescence, which can be abrogated in the case of PDK4 suppression (Fig. 4k-l). The results from fluorescence sensors suggest that the lactate production level increases in parallel to the NAD^+^/NADH ratio decrease in senescent cells, while both changes are correlated with PDK4 activity. The data not only disclose the concurrent fluctuation of NAD^+^/NADH conversion and lactate generation, but further substantiate the central role of PDK4 in orchestrating a metabolic profile specifically associated with the occurrence of cellular senescence.

### Stromal cells expressing PDK4 alter the expression profile and phenotypes of cancer cells *via* HTR2B

We next sought to determine the influence of stromal cells expressing PDK4 on their surrounding microenvironment, specifically cancer cells. Since PSC27 originally derived from human prostate, we first chose to examine PCa cells. PSC27-derived CM were employed to treat PCa cells in culture, with cancer cells subject to genome-wide analysis. Data from RNA sequencing (RNA-seq) indicated that 4188 transcripts were significantly upregulated or downregulated (fold change > 2, *P* < 0.05) in PC3 cells, with 4860 and 3756 transcripts changed in DU145 and M12 cells, respectively (Fig. 5a). We noticed remarkable and comprehensive changes in the biological processes of PCa cells, as evidenced by typically affected activities in signal transduction, cell communication, intracellular transport, energy pathways and metabolism regulation (Fig. 5b and Supplementary Fig. 5a, b). Together, the data suggest a salient capacity of PDK4-expressing stromal cells in reprogramming the transcriptome of recipient cancer cells through production of the CM, the latter of a typically acidic nature.

**Fig. 5.**
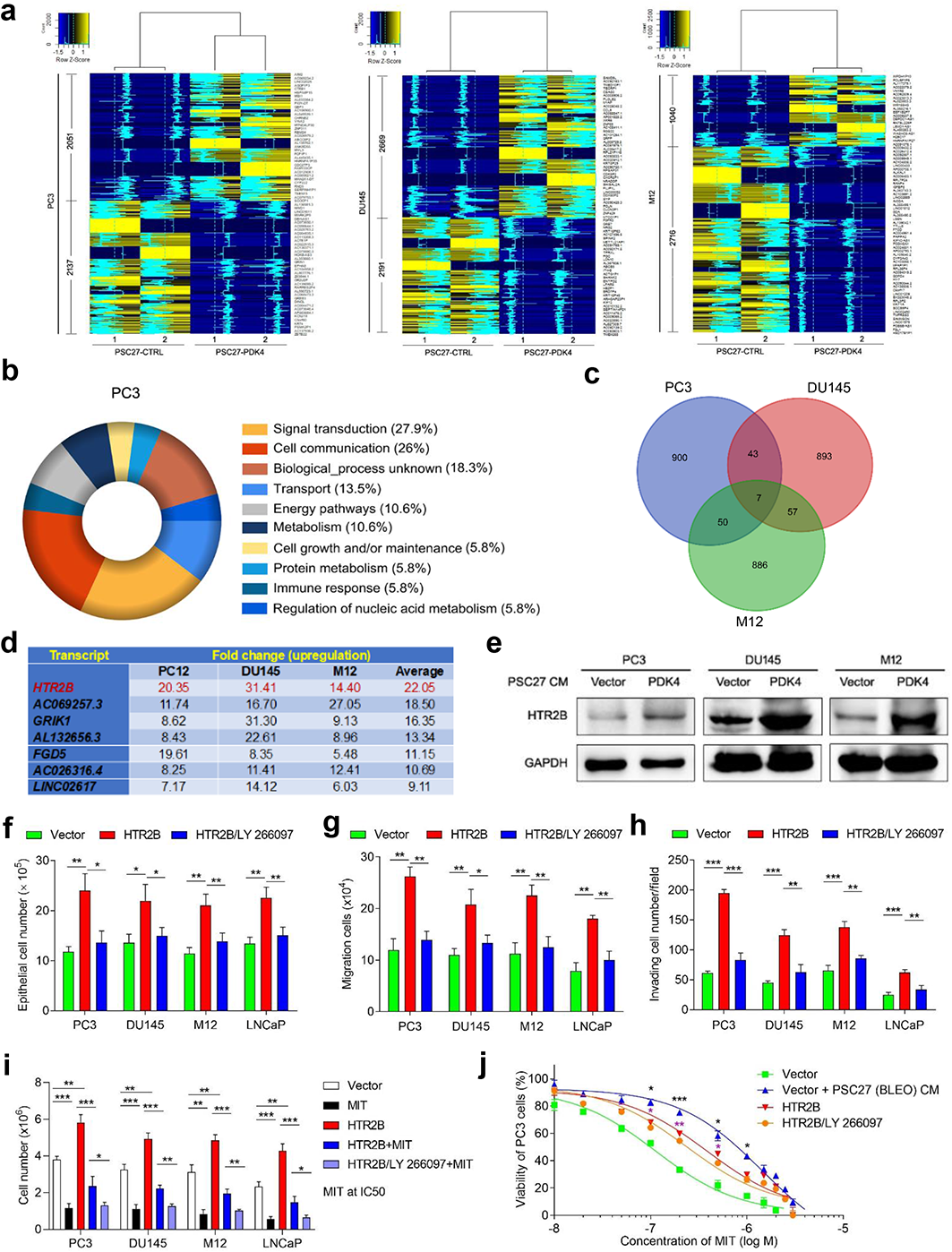
Stromal expression of PDK4 induces profound changes of PCa cell expression profile and causes enhanced malignancy. **a.** Heatmap depicting differentially expressed human transcripts in PCa lines including PC3, DU145 and M12 after a 3-d culture with the CM collected from PSC27 cells overexpressing PDK4 (PSC27-PDK4). In contrast to cancer cells cultured with control CM (PSC27-CTRL), the number of genes up- and downregulated *per* PCa line are indicated. The intensity of tracing lines are consistent with the relative expression fold change averaged *per* up- and downregulated genes. **b.** Graphical visualization of pathways by GO profiling (pie chart depicting biological processes). Those significantly enriched genes in the upregulated list were sorted according to their fold change in PC3 cells exposed to the CM of PSC27-PDK4 cells. **c.** Venn diagram displaying the overlap of transcripts co-upregulated in PC3, DU145 and M12 cells (*per* 2 or 3 lines) upon in culture treatment with the CM from PSC27-PDK4 stromal cells in contrast to those treated with the CM of PSC27-CTRL. **d.** Summary of transcripts co-upregulated in PC3, DU145 and M12 lines (top ranked, with a fold change ≥ 5.0 and FDR < 0.01) upon treatment with the CM of PSC27-PDK4 stromal cells. Red highlighted, HTR2B. **e.** Immunoblot assessment of protein expression of HTR2B in the three PCa lines. β-actin, loading control. **f.** Measurement of the proliferation capacity of PCa lines at different conditions. As experimental group, human HTR2B was transduced into each PCa line, with LY 266097 applied to media as a drug treatment control. **g.** Examination of the 2D migration activity of PCa lines at different conditions. Cells were treated in a manner similar to that described in (**f**). **h.** Evaluation of the invasion ability of PCa lines at different conditions. Cells were treated in a manner similar to that described in (**f**). **i.** Determination of the resistance of PCa lines to MIT upon exposure to the CM of PSC27-PDK4 stromal cells. MIT, mitoxantrone, a chemotherapeutic agent applied at the IC50 concentration *per* cell line established prior to the assay. **j.** Dose-response curves (non-linear regression/curve fit) plotted from MIT-based viability assays of PC3 exposed to the CM of PSC27-PDK4 cells and treated by a range of concentrations of MIT. Data in all bar plots are shown as mean ± SD. Data in **a**, **e-j** are representative of 3 independent experiments. All *P* values were calculated by Student’s *t*-tests. ^, *P* > 0.05. *, *P* < 0.05. **, *P* < 0.01. ***, *P* < 0.001.

Among the transcripts significantly upregulated by the CM of PSC27 cells (*P* < 0.05, FDR < 0.01, top 1000 shown *per* PCa line, Supplementary Table 1), we noticed that there were 7 transcripts showing up and commonly expressed by PC3, DU145 and M12 cells (fold change > 4, *P* < 0.01) (Fig. 5c, d). We then chose to focus on a specific gene (or more accurately, transcript), which potentially contributes to malignant alterations of cancer cells. Immunoblots substantiated that expression of 5-hydroxytryptamine receptor 2B (HTR2B), a gene encoding one of the several different receptors for serotonin which belongs to the G-protein coupled receptor 1 family, was markedly induced upon exposure of PCa cells to the CM of PDK4-expressing stromal cells (Fig. 5e). Serotonin is a biogenic hormone that functions as a neurotransmitter and a mitogen, while serotonin receptors mediate the central and peripheral physiologic functions of serotonin, including regulation of cardiovascular functions and impulsive behavior. We interrogated the implications of HTR2B in phenotypic changes of individual PCa cell lines. Of note, the capacities of proliferation, migration and invasion of PCa cells were comprehensively enhanced, as evidenced by our *in vitro* assays (Fig. 5f-h). More importantly, transduction of HTR2B enhanced the resistance of PCa cells to MIT, a DNA-targeting chemotherapeutic agent administered to cancer patients including those developing PCa ^38, 39^ (Fig. 5i). The survival curves of cancer cells under genotoxic stress of MIT displayed a shift toward higher concentrations of this drug, as exemplified by the case of PC3 (Fig. 5j). Notably, the CM derived from senescent stromal cells generated by BLEO treatment (PSC27 (BLEO) CM) caused a more dramatic increase of chemoresistance than HTR2B *per se*, suggesting the presence and contribution of other molecules in the CM of senescent stromal cells, particularly a large number of soluble factors encoded by the full-spectrum of the SASP. However, these gain of functions were generally lost when LY 266097 ^40^, a selective antagonist of HTR2B, was applied to the culture, suggesting that enhanced malignancies of cancer cells were mainly attributed to the activity of HTR2B after ectopically expressed in these lines (Fig. 5f-j).

As HTR2B is one of the most upregulated genes observed in PCa lines we examined upon treatment with PDK4^+^ stromal cell CM, whether or not it accounts for the principal force driving malignant changes of recipient cancer cells remain unknown. We then used LY 266097 or gene-specific small hairpin RNAs (shRNAs) to target HTR2B in individual PCa cell lines before performing phenotypic assays. Interesting, the gain of functions conferred by the CM of PDK4^+^ stromal cells was substantially abrogated in the absence of HTR2B or upon LY 266097 treatment (Supplementary Fig. 5c-g). Thus, HTR2B is a competent factor that mediates the influence of the acidic microenvironment generated by PDK4-expressing stromal cells on recipient cancer cells, while elimination of HTR2B or blockade of HTR2B signaling in cancer cells holds the potential to remarkably weaken their malignant phenotypes.

### Therapeutically targeting PDK4 improves chemotherapeutic outcome in preclinical trials

Given the acidic extracellular microenvironment formed by stromal cells expressing PDK4 and its effects of on the biological phenotype and expression profile of cancer cells *in vitro*, we were tempted to query the pathological consequences of PDK4 induction in the TME under *in vivo* conditions. To this end, we constructed tissue recombinants by admixing PSC27 sublines with PC3 cells at a pre-optimized ratio of 1:4 before subcutaneous implantation to the hind flank of experimental mice with severe combined immunodeficiency (SCID). The animals were gauged for tumor size at the end of an 8-week period. Compared with tumors comprising PC3 and PSC27^Vector^, xenografts composed of PC3 and PSC27^PDK4^ displayed significantly increased sizes (*P* < 0.01) (Supplementary Fig. 6a). Conversely, PDK4 knockdown (by shRNA) from these PSC27^PDK4^ cells prior to xenograft implantation markedly reduced tumor volumes (*P* < 0.01 and *P* < 0.05, respectively).

To closely mimic clinical conditions involving chemotherapeutic agents, we designed a preclinical regimen which incorporates a genotoxic drug (MIT) and/or the PDK4 inhibitor (PDK4-IN) (Fig. 6a). Two weeks after cell implantation when stable uptake of tumors by host animals were generally observed, a single dose of MIT or placebo was administered at the 1^st^ day of 3^rd^, 5^th^ and 7^th^ week until the end of the 8-week regimen (Supplementary Fig. 6b). Although PDK4-IN administration did not provide noticeable benefits, MIT treatment caused remarkable tumor shrinkage (57.5% volume reduction), validating the efficacy of MIT as a cytotoxic agent (Fig. 6b). Importantly, when PDK4-IN was combined with MIT, a further decline of tumor volume was observed (35.5%), resulting a total shrinkage by 72.6% as compared with the vehicle control (Fig. 7b).

**Fig. 6.**
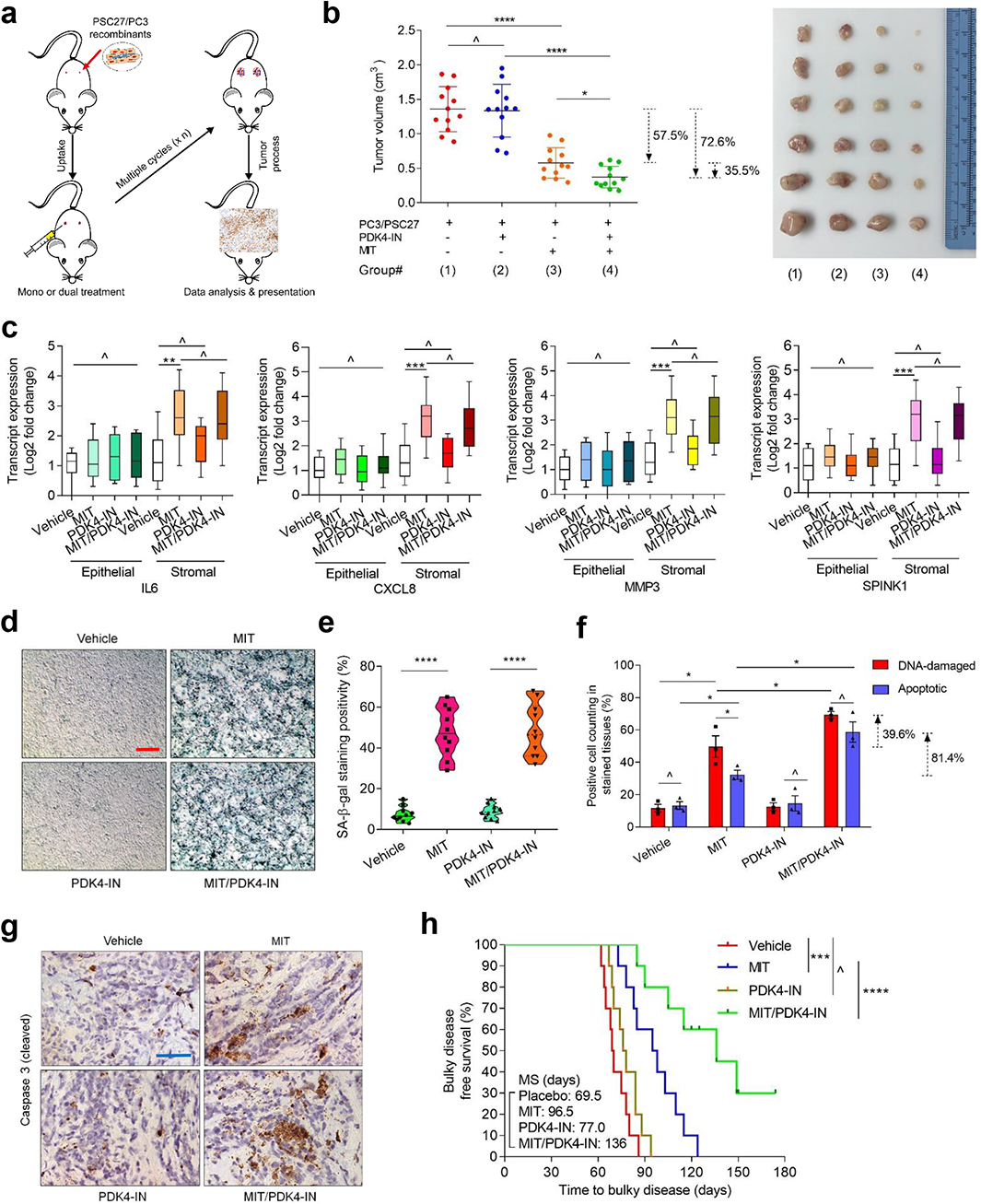
Therapeutically targeting PDK4 in the damaged TME promotes therapeutic outcome in preclinical trials. **a.** Schematic workflow of experimental procedure for severe combined immunodeficient (SCID) mice. Two weeks after subcutaneous implantation and *in vivo* uptake of tissue recombinants, animals received either single or combinational agents administered as metronomic treatments composed of several cycles. **b.** Statistical profiling of tumor end volumes. PC3 cells were xenograted alone or together with PSC27 cells to the hind flank of SCID mice. MIT and PDK4-IN was administered either alone or concurrently to induce tumor regression. Right, representative tumor images. **c.** Transcript assessment of several canonical SASP factors expressed in stromal cells isolated from the tumors of SCID mice. Tissues from animals implanted with both stromal and cancer cells were subject to LCM isolation, total RNA preparation and expression assays. **d.** Representative IHC images of SA-β-Gal staining profile of tissues isolated from placebo or drug-treated animals. Scale bar, 100 μm. **e.** Comparative statistics of SA-β-Gal staining for mouse tissues described in (**d**). **f.** Statistical assessment of DNA-damaged and apoptotic cells in the tumor specimens analyzed in (**f**). Values are presented as percentage of cells positively stained by IHC with antibodies against γ-H2AX or caspase 3 (cleaved). **g.** Representative IHC images of caspase 3 (cleaved) in tumors at the end of therapeutic regimens. Biopsies of placebo-treated animals served as negative controls for drug-treated mice. Scale bars, 100 μm. **h.** Bulky disease free survival plotted against the time of recombinant tissue injection until animal death attributed to the development of advanced bulky diseases. MS, median survival. *P* values were calculated by two-sided log-rank (Mantel–Cox) tests. Data in all dot or bar graphs are shown as mean□±□S.D. and representative of 3 independent experiments. All *P* values were calculated by Student’s *t*-tests. ^, *P* > 0.05. *, *P* < 0.05. **, *P* < 0.01. ***, *P* < 0.001. ****, *P* < 0.0001.

**Fig. 7.**
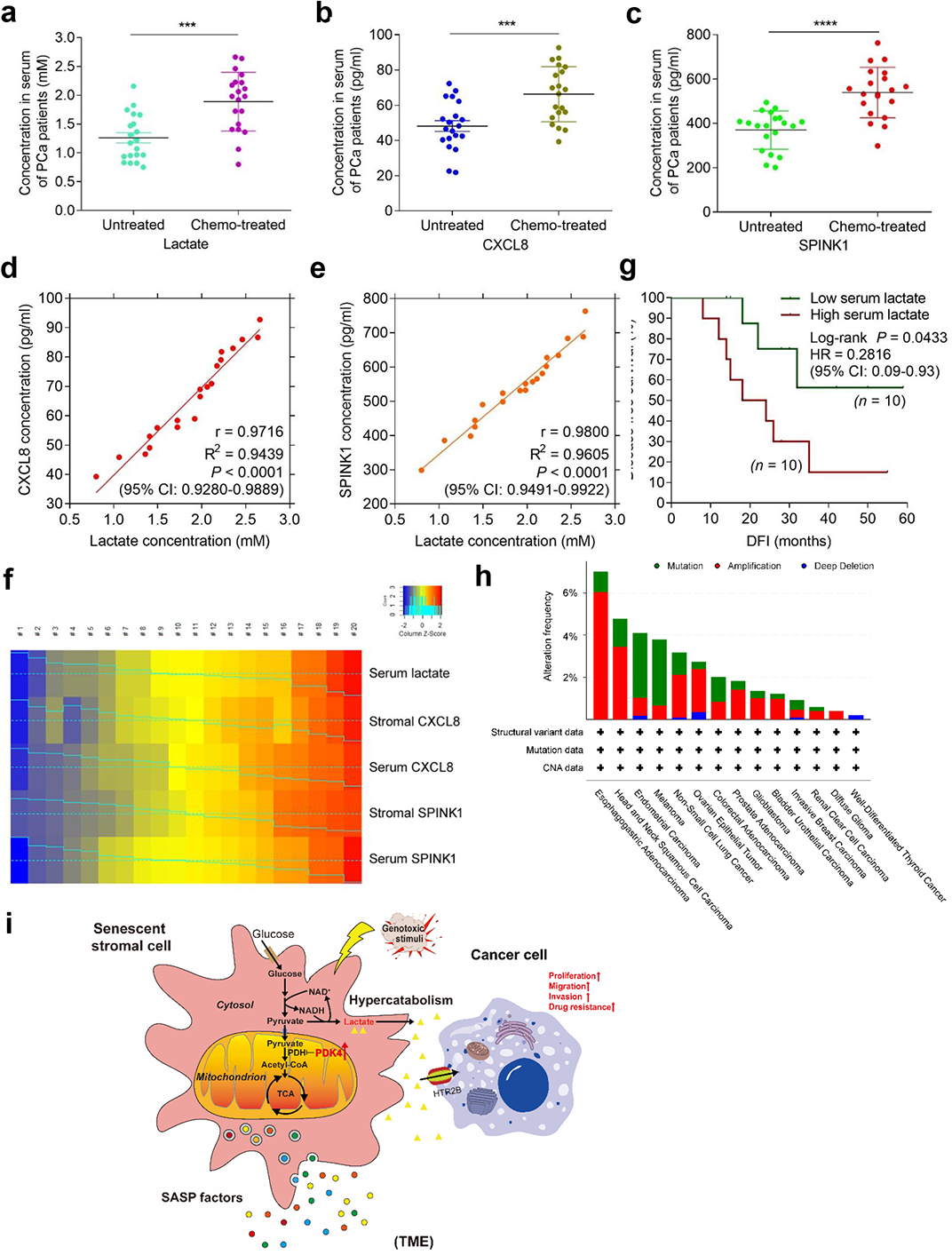
Lactate is a novel circulating biomarker indicative of the SASP *in vivo* and predicts adverse therapeutic outcome in cancer clinics. **a.** Abundance of lactate in the serum of untreated and chemo-treated PCa patients. Data were derived from ELISA measurement and shown as mean ± SD. N = 20. **b.** Abundance of CXCL8 protein in patient serum analyzed in (**a**). Data from ELISA assays and presented as mean ± SD. N = 20. **c.** Abundance of SPINK1 protein in patient serum analyzed in (**a**). Data from ELISA assays and presented as mean ± SD. N = 20. **d.** Scatterplot showing correlation between lactate and CXCL8 in the serum of individual patients described in (**a-c**). Pearson’s correlation coefficient, *P* value and confidence interval are indicated. **e.** Scatterplot showing correlation between lactate and SPINK1 in the serum of individual patients described in (**a-c**). Pearson’s correlation coefficient, *P* value and confidence interval are indicated. **f.** Heatmap depicting the overall correlation between serum lactate, stromal CXCL8, serum CXCL8, stromal SPINK1 and serum SPINK1 in chemo-treated patients (n = 10). The raw scores of stromal CXCL8 and SPINK1 were derived from independent pathological reading of primary tumor tissues of PCa patients, with those of serum CXCL8 and SPINK1 obtained from ELISA assays. Color key, relative expression of these factors in stromal tissue or patient serum. **g.** Kaplan-Meier survival analysis of chemo-treated PCa patients. Disease free survival (DFS) stratified according to circulating lactate in serum (low, average score < 2, dark green; high, average score ≥ 2, dark red). DFS represents the length (months) of period calculated from the date of chemotherapy to the point of first time disease relapse. Survival curves generated according to the Kaplan–Meier method, with *P* value calculated using a log-rank (Mantel-Cox) test. N = 10 per group. **h.** TCGA data showing alterations of PDK4 in a variety of human cancer types at the genomic level, including mutation, amplification and deep deletion. Alteration frequency is displayed in percentage. **i.** Graphic illustration to summarize metabolic reprogramming of senescent cells and formation of an acidic microenvironment in a genotoxic setting and functional implications of the metabolite lactate in promoting cancer resistance against chemotherapeutic treatments. Data in **a**-**e** are representative of 3 independent experiments. *P* values were calculated by Student’s *t*-test (**a**-**c**), Pearson test (**d**-**e**) and log-rank (Mantel-Cox) test (**g**). ***, *P* < 0.001. ****, *P* < 0.0001.

Not surprisingly, we observed a considerable upregulation of typical SASP factors such as IL6, CXCL8, MMP3, SPINK1 and AREG, accompanied by expression of typical senescence markers including p16^INK4a^ and p21^CIP1^ in stromal cells of xenografts composed of PC3/PSC27 cells, implying development of an *in vivo* senescence and expression of the SASP upon MIT treatment (Fig. 6c and Supplementary Fig. 6c). However, PDK4-IN neither induced nor affected cellular senescence and the SASP, as evidenced by results from its administration as a mono agent or combined with MIT (Fig. 6c and Supplementary Fig. 6c). Although senescence was induced in cancer cells in animals undergoing MIT treatment as suggested by p16^INK4a^ and p21^CIP1^ expression, we did not notice a typical and full-spectrum SASP in these cells of epithelial origin, results largely consistent with findings of our former studies ^41, 42^. Of note, PDK4 expression was remarkably induced in stromal cell populations, but not in their epithelial counterparts (Supplementary Fig. 6c), basically in line with our *in vitro* data (Fig. 1d, e). Histological staining indicated elevated SA-β-Gal positivity in tumor tissues of mice that experienced MIT or MIT/PDK4-IN treatment, confirming comprehensive senescence occurrence in these groups (Fig. 6d, e). In contrast, treatment with PDK4-IN itself did not seem to cause induction or suppression of senescence, thus congruent with the nature of this agent, which typically does not target DNA nor damage macromolecules (Fig. 6c-e).

Given the primary *in vivo* expression results derived from treatments involving MIT and/or PDK4-IN, we next asked how pharmacologically targeting PDK4 could enhance the therapeutic response of tumors. To disclose the possible mechanism(s), we chose to dissect tumors from animals 7 days after initiation of treatment, a timepoint right prior to the development of resistant colonies. In contrast to vehicle, MIT *per se* caused significant DNA damage and apoptosis in cancer cells (Fig. 6f). Although PDK4-IN alone did not cause typical DDR or cell apoptosis, it showed prominent efficacy in enhancing these therapeutic indices upon combination with MIT (*P* < 0.05). IHC staining disclosed increased caspase 3 cleavage, a canonical apoptosis indicator, upon MIT administration, a tendency that was further enhanced in the presence of PDK4-IN (Fig. 6g).

To expand, we used LNCaP, a second PCa cell line which expresses androgen receptor (AR) and is routinely employed as a hormone-responsive cell model. To produce an AR-naïve setting, we circumvented experimental castration, but followed the same protocol designed for PC3-tailored therapeutic cohorts. We noticed significantly reduced volumes of LNCaP/PSC27 tumors when mice underwent MIT/PDK4-IN co-treatment, in contrast to MIT administration only (36.1%) (Supplementary Fig. 6d). Similar results were observed when 22Rv1, a castration-resistant PCa cell line, was applied to replace LNCaP for *in vivo* assays (35.3%) (Supplementary Fig. 6e). As supporting efforts to exclude the possibility of tumor type specificity, we generated tumors composed of A549, a non-small cell lung cancer (NSCLC) line, and HFL1, a lung fibroblast line. The results largely produced those observed in animals carrying tumors developed from PCa lines (34.0%) (Supplementary Fig. 6f). Together, these data suggest that specific targeting of PDK4, a kinase responsible for lactate production during glucose metabolism and formation of an acidic microenvironment, specifically in a treatment-damaged TME which harbors a number of senescent cells, can substantially promote tumor regression in chemotherapeutic settings, a process independent of androgen regulation or AR signaling of prostate tumors *per se*. We hereby conclude that the resistance-minimizing effects of PDK4-targeting strategy are not limited to a specific cancer type, but may have implications to a wide range of malignancies.

We next assessed tumor progression consequence by comparing the survival of different animal groups in a time-extended preclinical cohort, with PCa mice as a pilot model. In the course of tumor growth monitoring, a bulky disease was considered developing once the tumor burden was prominent (size ≥ 2000 mm^3^), an approach described previously ^24, 43^. Mice that received MIT/PDK4-IN combinational treatment displayed the most prolonged median survival, gaining a 40.9% longer survival when compared with those treated by MIT only (Fig. 6h, green *vs* blue). However, PDK4-IN treatment alone did not achieve significant benefits, as it conferred only marginal survival advantage (Fig. 6h, brown *vs* red). Thus, targeting PDK4 in the TME affects neither tumor growth nor animal survival, while MIT/PDK4-IN co-treatment has the competence to significantly improve both parameters.

Importantly, to establish the safety and feasibility of such a therapeutic regimens, we conducted routine pathophysiological appraisal. The data supported that either single or combinatorial treatment was well tolerated, as evidenced by body weight maintenance throughout the therapeutic timeframe (Supplementary Fig. 7a). Further, there were no significant perturbations in the serum level of creatinine, urea and metabolic activities of liver enzymes (ALP and ALT) (Supplementary Fig. 7b). Additional data from mice developing NSCLC carcinomas and treated by DOX/PDK4-IN, or DOX/PDK4-IN-treated immunocompetent animals (C57BL/6J background), generally phenocopied PCa animals by manifesting no routine blood count fluctuations, thus further validating these findings (Supplementary Fig. 7c-f). Our data support that strategies combining a PDK4-targeting agent with classical chemotherapy hold the potential to enhance tumor responses without causing severe systemic cytotoxicity.

### TIS-associated serum lactate adversely predicts posttreatment survival of cancer patients

Although higher PDK4 expression in the tumor foci is correlated with lower survival of posttreatment in clinic settings (Fig. 2g and Supplementary Fig. 2d), whether the metabolic product lactate derived from the TME harboring stromal cells developing TIS is technically detectable and can serve as a marker for clinic purposes remains largely unclear. To address this, we acquired peripheral blood samples from PCa patients, including one cohort that experienced standard chemotherapy and the other that did not. ELISA assays of the serum from chemo-treated patients revealed that lactate levels in the treated cohort were significantly higher than that of the treatment-naïve group (Fig. 7a). The pattern was essentially reproduced by a remarkable increase of CXCL8 and SPINK1, canonical hallmarks of the SASP, in the same cohort of posttreatment patients (Fig. 7b, c). The data suggest that a circulating scale of lactate, the product of glucose metabolism *via* the glycolysis branch, emerges in the peripheral blood alongside with an *in vivo* SASP, which is intimately correlated with *in situ* senescence of tissues and develops after chemotherapeutic regimens, and both are systemically traceable in the serum of cancer patients. Further, it is intriguing to determine whether the blood level of lactate is correlated with that of typical SASP factors such as CXCL8 and SPINK1 in the same individual patients after clinical treatment. Subsequently analysis of ELISA data disclosed a significant and positive correlation between lactate and CXCL8, as well as between lactate and SPINK1 (Fig. 7d, e). Thus, lactate production and SASP expression is mutually linked, largely resembling the close correlation between PDK4 induction and the SASP factors as revealed by our clinical data derived from tumor samples *per se* (Fig. 2f).

We then expanded the study by longitudinal analysis of these factors in both primary tumor foci and peripheral blood (20 chemo-treated patients randomly selected). Surprisingly, cross-organ comparisons indicated a pronounced association between in-tissue expression and circulating level *per* factor, with the amounts of lactate, CXCL8 and SPINK1 apparently varying in parallel either within the primary tissue or through peripheral blood of each individual (Fig. 7f). Together, our data suggests that lactate indeed represents one of the critical TME-derived biological factors precisely imaging the development of an *in vivo* SASP, and can be exploited to assess the SASP magnitude in posttreatment cancer patients.

Clinical profiling subsequently uncovered a negative correlation between plasma level of lactate and posttreatment survival of PCa patients, further substantiating the pathological impact of lactate, which as a TME-derived molecule directly predicts adverse outcome once the TME is subject to irreparable damage by clinical agents (Fig. 7g). As PDK4 is subject to frequent mutation, amplification and deep deletion as disclosed by the TCGA pan-cancer atlas studies (Querying 22179 patients/22802 samples in 36 clinical studies) which document global genomics data from multiple cancer types ^44, 45^ (Fig. 7h), this molecule represents an important predictor of disease progression in treatment-naïve patients in clinical oncology ^46, 47^. Contrasting former studies which mainly focus on the genomic alterations and pathological behaviors of cancer cells, we herein propose that routine surveillance of lactate, a major metabolic product derived from enhanced glycolysis driven by PDK4 highly expressed in stromal cells particularly those in the case of TIS, *via* a noninvasive avenue such as liquid biopsy, can provide a novel, practical and accurate strategy for both prognosis and prevention of advanced pathologies in clinical oncology.

## Discussion

Aging is defined as a complex and time-dependent process that causes a progressive decline of physiological integrity, particularly functional degeneration of multiple types of tissues and organs. A number of hallmarks contribute to aging, while cellular senescence is identified as one of the primary risk factors for the initiation and development of age-related conditions, such as cancers, diabetes, cardiovascular disorders and neurodegenerative diseases ^4, 48^. Senescent cells synthesize a large array of pro-inflammatory cytokines, chemokines and extracellular matrix degrading enzymes, a feature known as the SASP ^5^. Importantly, discovery of the SASP proposes a reasonable and critical mechanism to explain why senescent cells, even accumulating in a low number *in vivo* during aging, can generate large detrimental effects on organismal health. However, whether or not the SASP is the sole source of senescence-associated factors that contribute to the loss of tissue homeostasis and organ function, pathophysiological events directly linked with aging and age-related diseases, remains yet unknown. In this study, we mapped the metabolic landscape of glucose metabolism upon cellular senescence, and discovered that senescent cells develop a substantially reprogrammed metabolism and produce an increased amount of acidic metabolites, particularly the glycolysis product lactate, the latter mediated by PDK4 upregulation and potentiating the formation of an acidic microenvironment. With experimental models, we demonstrated that consequences of such a metabolic rewiring include but are not limited to increased cancer malignancy, specifically enhanced drug resistance, an event responsible for accelerated disease progression and reduced survival in the posttreatment stage of cancer individuals.

Reprogrammed metabolism is indeed one of the established hallmarks of cancer, while a distinctive abnormality of cancer cell metabolism is the conversion of glucose into lactate even upon normal oxygen supply, a phenomenon termed ‘Warburg effect’ ^49^. Despite a reduced efficiency of ATP production as compared with thorough catabolism of glucose through mitochondrial TCA cycle, aerobic glycolysis provides cancer cells with the biosynthetic advantage of fast uptake and incorporation of nutrients into biomass, which is essential for malignant expansion ^50^. Some key enzymes involved in aerobic glycolysis display aberrant activities in cancer cells, while others are subjected to genetic mutations and expression regulation in levels of transcription, translation and posttranslational modifications ^50, 51^. Metabolic switch from OXPHO to glycolysis in cancer cells regulates the invasion-metastasis cascade by promoting epithelial-mesenchymal transition (EMT), tumor angiogenesis and distant metastasis. In contrast to cancer cells, the vast majority of senescent cells have a remarkably lower rate of genetic mutations, mainly due to the arrested cell cycle or terminated proliferation. However, senescent cells are metabolically active, and display significant changes in mitochondrial dynamics and cytoplasmic organization ^28, 29^. The mitochondria of senescent cells can be elongated, enlarged and/or hyperfused, depending on different models of cellular senescence employed ^32^. We observed damaged ultrastructure of mitochondria upon cellular senescence, especially after treatment with genotoxic agents, an event suggestive of potential mitochondrial dysfunctions associated with oxidative stress. Further, as senescent cell are inherently resistant to programmed cell death, the hyperfused mitochondrial network we observed during senescence could be a protective mechanism underlying prolonged survival, or more specifically, apoptosis resistance ^52^.

PDK4 plays a pleiotropic, dynamic and context-dependent role in regulation of glucose and fatty acid metabolism. Pyruvate enters the TCA cycle through PDH, the gatekeeping enzyme with its activity regulated *via* reversible phosphorylation for entry of pyruvate into TCA cycle by controlling the conversion of pyruvate into acetyl-CoA, while PDKs suppress PDH activity and promotes a switch from mitochondrial oxidation to cytoplasmic glycolysis ^47^. Belonging to the PDK superfamily composed of 4 kinases in humans and rodents (PDK1-4) and acting as a nutrient sensor and critical regulator of glucose homeostasis, PDK4 has become an attractive target for treatment of various metabolic pathologies including hyperglycemia, insulin resistance and hepatic steatosis ^53^. Upregulation of PDK4 mediates aerobic glycolysis (Warburg effect), favors tumor growth and promotes apoptosis resistance ^20, 54–56^. However, potential implications of PDK4 in senescence-associated phenotypes remain hitherto underexplored. To this end, we made efforts to establish PDK4 expression pattern upon cellular senescence, elucidate its role in diverging glucose metabolism towards glycolysis to produce lactate, and unravel the correlation of PDK4 upregulation in tumor stroma and patient survival post-chemotherapy. Notably, PDK4 upregulation is responsible for increased production of lactate, a molecule that accumulates in the treatment-damaged TME but ultimately enters the circulating system. Of clinical relevance, PDK4 expression in the TME is significantly and negatively correlated with disease-free survival of cancer patients in the posttreatment stage.

Senescent cells can be less efficient in ATP production, generating a bioenergetic imbalance with an increased AMP/ATP ratio, a fact that could be partially explained by a decreased efficiency of OXPHOS (characterized by less H^+^ in the intermembrane space) associated with a reduction of mitochondrial membrane potential (MMP) ^57^. Upon cellular senescence, however, we observed enhanced mitochondrial respiration, ATP production and increased oxidative stress as evidenced by elevated ROS, the latter may be due to mitochondrial dysfunction *per se*. These changes are presumably correlated with the enhanced uptake flux of glucose in senescent cells, which supports to meet the increased needs of energy and biosynthesis materials when the intracellular protein machinery is functionally engaged in synthesis of the full-spectrum SASP factors. TIS cells developing the SASP display endoplasmic reticulum stress, an unfolded protein response (UPR) and enhanced ubiquitination, while TIS cancer cells are more sensitive to glucose utilization blockade, a baseline for their selective elimination through pharmacological targeting of the metabolic demands ^28^. Given our findings that unmask the hypercatabolic nature of senescent cells such as those undergoing TIS, it is reasonable to speculate the potential of therapeutically targeting metabolic features of these cells, a new modality of senotherapy exploitable to intervene aging and age-related conditions.

Despite an increased number of mitochondria, senescent cells experience a shift towards glycolysis, a phenomenon previously reported in replicatively senescent cells, which exhibit increases in both glucose uptake and lactate production prior to morphological alterations characteristic of senescence ^58^. Similar findings were derived from more sophisticated assays including metabolomics or measurements of mitochondrial oxygen flux and ECARs in different senescent cell types ^59, 60^. However, such a metabolic shift seems to be not universal for all types of cells and senescent inducers. For example, in the case of OIS, glucose uptake was not changed, but oxygen consumption and TCA activity were increased, changes linked to elevated PDH activity ^29^. Interestingly, enhanced glycolysis *via* overexpression of glycolytic enzymes phosphoglycerate mutase or glucosephosphate isomerase allows cells to bypass both replicative and OIS ^61^. Replicative senescent human mammary epithelial cells exhibit neither an overall glycolytic shift, glucose consumption nor lactate secretion ^57^. In contrast, our metabolic data suggest that senescent human stromal cells in the case of TIS, more specifically, genotoxicity-induced senescence (GIS), develop a rewired metabolism, with increased glucose utilization, elevated TCA activity, augmented ATP production and enhanced lactate production, further proposing the caveat to avoid overgeneralizations for senescence-associated metabolism, which does vary according to the specific cell type and/or physiological context.

As key cofactors for many metabolic reactions, such as glycolysis, the TCA cycle and fatty acid β-oxidation, NAD^+^ and its reduced form NADH play central roles in mammalian energy metabolism, while the conversion between these two forms drives the production of ATP *via* anaerobic glycolysis and mitochondrial OXPHO ^62^. NAD^+^ is also a co-substrate of many regulatory enzymes, including sirtuins, CD38 and PARPs, directly and indirectly influencing many key cellular functions including DNA repair, chromatin remodeling and cellular senescence ^13^. The SoNar sensor is unique as its fluorescence sensitively responds to the binding of NAD^+^ or NADH and the read-out reflects the NAD^+^/NADH ratio rather than the absolute concentrations of the two nucleotides intracellularly, thus serving a powerful tool for NAD^+^/NADH detection and bioimaging. In this study, we observed a significantly decreased NAD^+^/NADH ratio as revealed by the SoNar sensor, a trend indicative of markedly increased glycolytic activity and enhanced mitochondrial OXPHO, events correlated with an active conversion from NAD^+^ towards NADH. Importantly, such a remarkable change in NAD^+^/NADH ratio is accompanied by a significant increase of cytosolic lactate level in senescent cells. The capacity of lactate production is usually maintained homeostatically for proper functioning of organisms, while lactate accumulation in the TME can result in transcriptional alterations of genes correlated with cell fate regulation and phenotypic modulation ^63^. We noticed a significant enhancement of lactate production upon cellular senescence, as indicated by FiLa sensor, a case reflecting intracellular lactate accumulation under our experimental condition. As expected, the FiLa sensor responds well to lactate fluctuations in different contexts, providing a unique yet powerful tool for monitoring live-cell lactate levels, which is physiologically, or in certain settings, pathophysiologically relevant as a key metabolic indicator or regulator of energy metabolism. Of note, changes of both NAD^+^/NADH metabolism and lactate production appeared to be intimately correlated with PDK4 activity, further substantiating the critical role of this enzyme in shaping the activity of senescent cells, which exhibit a metabolic spectrum markedly distinct from that of their proliferating counterparts.

Metabolism plays an important role in regulating cellular senescence, a process that dramatically affects the aging process ^64^. By metabolic profiling and functional assessments of the glucose consumption axis, we hereby mapped the metabolic landscape of human senescent cells, and propose that PDK4, a PDH-modifying enzyme, acts as a key regulator of biochemical activities to shape a unique form of metabolism, namely hypercatabolism, upon cellular senescence. The observation that PDK4 activity is induced during senescence and plays a critical role in forming an acidic microenvironment underscores this enzyme as a valuable target to counteract the overall pathological impact of senescent cells. As of note, acidic microenvironments bridge the gap between high lactate levels, chronic inflammation and pathological events, while accumulation of lactate in solid organs is a pivotal and early event in various diseases ^65, 66^. Given that high PDK4 expression causes PDH phosphorylation, rewires energy metabolism, holds the potential to drive tissue homeostasis imbalance and physical dysfunction, our study raises the possibility that this metabolic kinase is pharmacologically exploited for therapeutic intervention of aging and multiple age-related disorders, including but not limited to cancer.

## Methods

### Cell culture

Primary normal human prostate stromal cell line PSC27 and breast stromal cell line HBF1203 were generously provided by Dr. Peter Nelson (Fred Hutchinson Cancer Research Center) and maintained in stromal complete medium as described previously ^23^. Human fetal lung stromal lines WI38 and HFL1, and foreskin stromal line BJ were from ATCC and cultured with F-12K medium supplemented with 10% FBS. Prostate cancer epithelial cell lines PC3, DU145, LNCaP and VCaP (ATCC) were routinely cultured with RPMI 1640 (10% FBS). Prostate cancer epithelial line M12 was a kind gift from Dr. Stephen Plymate (University of Washington), which originally derived from the benign line BPH1 but phenotypically neoplastic and metastatic ^67^. All lines were routinely tested for mycoplasma contamination and authenticated with STR assays.

### Vectors, viruses and infection

Full length human p16^INK4a^, HRas^G12V^ and PDK4 sequences was cloned into pLenti-CMV/To-Puro-DEST2 (Invitrogen), individually, as described ^23^. Small hairpin RNAs (shRNA) targeting sequences for specific genes were cloned in pLKO.1-Puro vector (Addgene). Upon production by 293T cells, lentivirual titers were adjusted to infect ∼ 90% of cells. Stromal cells were infected overnight in the presence of polybrene (8 μg/ml), allowed to recover for 48 h and selected for 72 h before used for further analysis. For expression of target genes in either stromal or epithelial cells, total RNA was prepared and subject to qRT-PCR assays (primers listed in Supplementary Table 2, Supporting Information).

### Cell treatments

Stromal cells were grown until 80% confluent (CTRL) and treated with 50 μg/ml bleomycin (BLEO), 5 μM doxorubicin (DOX), 2 μM mitoxantrone (MIT), 50 nM docetaxel (DTX), 50 μM paclitaxel (PTX) or 20 μM vinblastine (VBL). After treatment, the cells were rinsed briefly with PBS and allowed to stay for 7-10 days prior to performance of various examinations.

### Human cancer patient recruitment and biospecimen analysis

Administration of chemotherapeutic agents was performed for primary PCa patients (Clinical trial no. NCT03258320) and infiltrating ductal BCa patients (NCT02897700), by following the CONSORT 2010 Statement (updated guidelines for reporting parallel group randomized trials). Patients with a clinical stage ≥ I subtype A (IA) (T1a, N0, M0) of primary cancer but without manifest distant metastasis were enrolled into the multicentred, randomized, double-blinded and controlled pilot studies. Age between 40-75 years with histologically proven PCa, or age ≥ 18 years with histologically proven infiltrating ductal BCa was required for recruitment into the clinical cohorts. Data regarding tumor size, histologic type, tumor penetration, lymph node metastasis, and TNM stage were obtained from the pathologic records. Tumors were processed as FFPE biospecimens and sectioned for histological assessment, with alternatively prepared OCT-frozen chunks processed *via* lase capture microdissection (LCM) for gene expression analysis. Specifically, stromal compartments associated with glands and adjacent to cancer epithelium were separately isolated from tumor biopsies before and after chemotherapy using an Arcturus (Veritas Microdissection) laser capture microscope following previously defined criteria ^23^. The immunoreactive scoring (IRS) gives a range of 1-4 qualitative scores according to staining intensity *per* tissue sample. Categories for the IRS include 0-1 (negative), 1-2 (weak), 2-3 (moderate), 3-4 (strong) ^68^. The diagnosis of PCa and BCa tissues was confirmed based on histological evaluation by independent pathologists. Randomized control trial (RCT) protocols and all experimental procedures were approved by the Institutional Review Board of Shanghai Jiao Tong University School of Medicine, with methods carried out in accordance with the official guidelines. Informed consent was obtained from all subjects and the experiments conformed to the principles set out in the WMA Declaration of Helsinki and the Department of Health and Human Services Belmont Report.

### Histology and immunohistochemistry

Formalin-fixed paraffin-embedded (FFPE) tissue sections of 7-10 μm were deparaffinized in xylenes and rehydrated through a graded series of alcohols. Routine histology appraisal was performed with hematoxylin and eosin staining. For immunohistochemical (IHC) evaluation, FFPE sections experienced antigen retrieval with sodium citrate, incubation with 3% H_2_O_2_, treatment with avidin/biotin blocking buffer (Vector Laboratories) and then 3% BSA for 30 min. Staining with primary and secondary antibodies was conducted at 4°C for overnight and at room temperature for 60 min, respectively. Sections were incubated with a H_2_O_2_-diaminobenzidine (DAB) substrate kit (Vector, SK-4100). Samples were counterstained with hematoxylin, dehydrated and mounted. IHC images were obtained using an upright microscope (Olympus BX51). Brown staining indicated the immunoreactivity of samples.

### RNA-seq and bioinformatics analysis

Total RNA samples were obtained from PC3 and DU145 cells cultured with CM of either PSC27^Vector^ or PSC27^PDK4^. Sample quality was validated by Bioanalyzer 2100 (Agilent), and RNA was subjected to sequencing by Illumina NovaSeq 6000 with gene expression levels quantified by the software package RSEM (https://deweylab.github.io/RSEM/). Briefly, rRNAs in the RNA samples were eliminated using the RiboMinus Eukaryote kit (Qiagen, Valencia, CA, USA), and strand-specific RNA-seq libraries were constructed using the TruSeq Stranded Total RNA preparation kits (Illumina, San Diego, CA, USA) according to the manufacturer’s instructions before deep sequencing.

Pair-end transcriptomic reads were mapped to the reference genome (GRCh38.p13) (http://asia.ensembl.org/Homo_sapiens/Info/Index) (ensembl_105) with reference annotation from Gencode v27 using the Bowtie tool. Duplicate reads were identified using the picard tools (1.98) script mark duplicates (https://github.com/broadinstitute/picard) and only non-duplicate reads were retained. Reference splice junctions are provided by a reference transcriptome (Ensembl build 73)^69^. FPKM values were calculated using Cufflinks, with differential gene expression called by the Cuffdiff maximum-likelihood estimate function ^70^. Genes of significantly changed expression were defined by a false discovery rate (FDR)-corrected *P* value < 0.05. Only ensembl genes 73 of status “known” and biotype “coding” were used for downstream analysis.

Reads were trimmed using Trim Galore (v0.3.0) (http://www.bioinformatics.babraham.ac.uk/projects/trim_galore/) and quality assessed using FastQC (v0.10.0) (http://www.bioinformatics.bbsrc.ac.uk/projects/fastqc/). Differentially expressed genes were subsequently analyzed for enrichment of biological themes using the DAVID bioinformatics platform (https://david.ncifcrf.gov/), the Ingenuity Pathways Analysis program (http://www.ingenuity.com/index.html). Raw data of RNA-seq were deposited in the NCBI Gene Expression Omnibus (GEO) database under the accession code GSE198110.

### Venn diagrams

Venn diagrams and associated empirical *P*-values were generated using the USeq (v7.1.2) tool IntersectLists ^71^. The t-value used was 22,008, as the total number of genes of status “known” and biotype “coding” in ensembl genes 73. The number of iterations used was 1,000.

### RNA-seq heatmaps

For each gene, the FPKM value was calculated based on aligned reads, using Cufflinks ^70^. Z-scores were generated from FPKMs. Hierarchical clustering was performed using the R package heatmap.2 and the distfun = “pearson” and hclustfun = “average”.

#### Immunoblot and immunofluorescence analysis

Whole cell lysates were prepared using RIPA lysis buffer supplemented with protease/phosphatase inhibitor cocktail (Biomake). Nitrocellulose membranes were incubated overnight at 4°C with primary antibodies listed in Table S6, Supporting Information, and HRP-conjugated goat anti-mouse or -rabbit served as secondary antibodies (Vazyme). For immunofluorescence analysis, cells were fixed with 4% formaldehyde and permeabilized before incubation with primary and secondary antibodies, each for 1 hr. Upon counterstaining with DAPI (0.5 μg/ml), samples were examined with an Imager.A1 (Zeiss) upright microscope to analyze specific gene expression. Uncropped scans of the most important blots are provided in Figure S9, Supporting Information.

#### *In vitro* cell phenotypic characterization

For proliferation assays of cancer cells, 2 × 10^4^ cells were dispensed into 6 well-plates and co-cultured with conditioned medium (CM) from stromal cells. Three days later, cells were digested and counted with hemacytometer. For migration assays, cells were added to the top chambers of transwells (8 μm pore), while stromal CM were given to the bottom. Migrating cells in the bottom chambers were stained by DAPI 12-24 hr later, with samples examined with Axio Observer A1 (Zeiss). Invasion assays were performed similarly with migration experiments, except that transwells were coated with basement membrane matrix (phenol red free, Corning). Alternatively, cancer cells were subject to wound healing assays conducted with 6-well plates, with healing patterns graphed with bright field microscope. For chemoresistance assays, cancer cells were incubated with stromal CM, with the chemotherapeutic agent MIT provided in wells for 3 days at each cell line’s IC50, a value experimentally predetermined. Cell viability was assayed by a CCK8 kit, with the absorbance at 450 nm measured using a microplate reader.

#### Metabolic analysis

Extracellular acidification rate (ECAR) was measured with a Glycolysis Stress Test kit (Agilent Technologies, 103020-100), while oxygen consumption rate (OCR) assessed using a Cell Mito Stress Test kit (Agilent Technologies, 103015-100). ECAR and OCR were determined with an XF24 Extracellular Flux Analyzer (Seahorse Bioscience, North Billerica, MA, 01862) according to the manufacturer’s standard protocol. PSC27 was seeded at a density of 5 × 10^4^ cells/ well in the XF24 cell culture microplate (Agilent Technologies, 04721 and Q01321) at a 37[5% CO_2_ incubator for overnight. To measure ECAR, 10 mM glucose, 1 μM oligomycin and 50 mM 2-DG were injected into each well. To measure the OCR, 1.5 μM oligomycin, 0.5 μM carbonyl cyanide 4-(trifluoromethoxy) phenylhydrazone (FCCP) and 0.5 μM rotenone/antimycin were injected sequentially in order into each well. All Seahorse data were normalized with cell numbers. All metabolic parameters are automatically calculated by WAVE software equipped in the Seahorse. Each value was calculated as follows. Non-glycolytic acidification was referred to as last rate measurement prior to glucose injection. Glycolysis rate was referred to as maximum rate measurement before Oligomycin injection - last rate measurement before glucose injection. Glycolytic capacity was referred to as maximum rate measurement after Oligomycin injection - last rate measurement before glucose injection. For the OCR, basal respiration was referred to as last rate measurement before first injection - minimum rate measurement after Rotenone/Antimycin injection. ATP production was referred to as last rate measurement before Oligomycin injection - minimum rate measurement after Oligomycin injection.

#### Metabolite labelling and measurement by GC-MS

Cells were resuspended in 0.6 ml cold (−40°C) 50% aqueous methanol containing 100 µM norvaline as an internal standard, inserted in dry ice for 30 mins for thawing. Samples were added with 0.4 ml chloroform and vortexed for 30 second before centrifugation at 14,000 rpm (4 °C) for 10 min, with supernatant transferred to new 1.5 ml tubes for evaporation before stored at −80°C. Metabolites were processed for GC/MS analysis as follows: First, 70 μl of pyridine was added to the dried pellet and incubated for 20 min at 80 °C. After cooling, 30 µl of *N-tert*-butyldimethylsilyl-*N*-methyltrifluoroacetamide (Sigma) was added, with samples re-incubated for 60 min at 80 °C before centrifugation for 10 min at 14,000 rpm (4 °C). The supernatant was transferred to an autosampler vial for GC-MS analysis. A Shimadzu QP-2010 Ultra gas chromatography-mass spectrometry (GC-MS) was programmed with an injection temperature of 250°C and injected with 1 µl samples. GC oven temperature started at 110 °C for 4 min, before raised to 230 °C at 3 °C/min and to 280 °C at 20 °C /min with a final hold at this temperature for 2 min. GC flow rate with helium carrier gas was 50 cm/s, with the GC column used at 20 m x 0.25 mm x 0.25 mm Rxi-5ms. GC-MS interface temperature was 300°C, while ion source temperature (electron impact) was set at 200 °C with 70 V ionization voltage. The mass spectrometer was set to scan m/z range 50-800 with 1 kV detector.

GC-MS data were analyzed to determine isotope labeling. To determine ^13^C labeling, the mass distribution for known fragments of metabolites was extracted from the appropriate chromatographic peak. These fragments contained either the whole carbon skeleton of the metabolite, or lacked the alpha carboxyl carbon, or (for some amino acids) contained only the backbone minus the side-chain. For each fragment, the retrieved data comprised mass intensities for the lightest isotopomer (without any heavy isotopes, M0) and isotopomers with increasing unit mass (M1 to M6) relative to M0. These mass distributions were normalized by dividing by the sum of M0 to M6 and corrected for the natural abundance of heavy isotopes of the elements H, N, O, Si and C, using matrix-based probabilistic methods and implemented in MATLAB. Labeling results are expressed as average fraction the particular compound containing isotopic label from the particular precursor.

#### Lactate assay

Lactate production was measured using a Lactate Colorimetric Assay Kit (Sigma-Aldrich, MAK058). Cells were homogenized in lactate assay buffer and centrifuged at 13,000 x g for 10 min to remove insoluble materials. The supernatants were de-proteinized with a 10 kDa MWCO spin filter to remove other enzymes. Next, 50 ml of the supernatants was mixed with 50 ml of the reaction mix, with the reaction incubated for 30 min at room temperature. Lactate levels were measured at 450 nm using a microplate reader, with the relative level of lactate in all groups calculated and normalized to protein concentration.

#### Glucose uptake assay

The level of glucose uptake was measured using a Glucose Uptake-Glo Assay Kit (Promega, J1341), which provided a homogeneous bioluminescent method for assessing glucose uptake in mammalian cells based on the detection of 2-deoxyglucose-6-phosphate (2DG-6-P). Cells were removed from medium and then washed with PBS, afterwards 50 ml of 1mM 2-deoxyglucose (2DG) was added to the cells and incubated for 10 min at room temperature. Then 25 ml of acid detergent solution (stop buffer) was added to lyse the cells and terminate the uptake; 25 ml of high-pH buffer solution (neutralization buffer) was then added to neutralize the acid. Finally, 100 ml of 2DG-6-P detection reagent was added to the sample wells, with the reaction incubated at room temperature for 1-2 h. A Cytation 5 Cell Imaging Multi-Mode Reader was used to assay the luminescence. The relative level of glucose uptake in all groups was calculated and normalized to protein concentration.

#### Characterization of SoNar and FiLa *in vitro*

Purified protein was stored at −80 °C before experimental assays. For *in vitro* measurements, the purified sensor protein was diluted with 100 mM HEPES buffer containing 100 mM NaCl (pH 7.4). Fluorescence spectroscopy was performed on a fluorescence spectrophotometer (PerkinElmer, FL6500). Excitation spectra were recorded at an emission wavelength of 530 nm. Slit width was set as 10 nm bandpass and the PMT voltage was set at 500 V.

For nucleotide titration of SoNar and lactate titration of FiLa, the sensor protein was diluted in HEPES buffer (pH 7.4) to a final concentration of 0.2 μM. The fluorescence intensity was measured by a filter-based Synergy Neo 2 Multi-Mode microplate reader using 420 BP 20 nm or 485 BP 20 nm excitation and 532 BP 40 nm emission band-pass filters (BioTek). All solutions were prepared in HEPES buffer (pH 7.4). Each assay was performed in a 96-well black bottom plate using 50 μl of substrate and 50 μl of sensor protein. Fluorescence intensity was measured immediately.

#### Live-cell fluorescence imaging

For fluorescence imaging, normal and senescent (TIS) PSC27 cells stably expressing SoNar, FiLa, iNapc or FiLa-C were plated on 35 mm 4-chamber glass-bottom dish. The dosing group was treated with or without 5 μM PDK inhibitor for 1 h. Fluorescence images were acquired using a Leica TCS SP8 SMD confocal laser-scanning microscope system with HC Plan Apo CS2 63×1.40 NA oil objective. For dual-excitation ratio imaging, 405 nm excitation laser and 488 nm excitation laser with an emission range of 500-550 nm were used. Raw data were exported to ImageJ software as 12-bit TIF for analysis. The pixel-by-pixel ratio of the 405 nm excitation image by the 488 nm excitation image of the same cell was used to pseudo-color the images in HSB color space as previously described ^37, 72^.

#### Experimental animals and preclinical studies

All animals were maintained in a specific pathogen-free (SPF) facility, with NOD/SCID (Charles River and Nanjing Biomedical Research Institute of Nanjing University) mice at an age of approximately 6 weeks (∼20 g body weight) used. Ten mice were incorporated in each group, and xenografts were subcutaneously generated at the hind flank upon anesthesia mediated by isoflurane inhalation. Stromal cells (PSC27 or HBF1203) were mixed with cancer cells (PC3, LNCaP or MDA-MB-231) at a ratio of 1:4 (i.e., 250,000 stromal cells admixed with 1,000,000 cancer cells to make tissue recombinants before implantation *in vivo*). Animals were sacrificed at 2-8 weeks after tumor xenografting, according to tumor burden or experimental requirements. Tumor growth was monitored weekly, with tumor volume (v) measured and calculated according to the tumor length (l), width (w) and height (h) by the formula: v = (π/6) × ((l+w+h)/3)^3 41^. Freshly dissected tumors were either snap-frozen or fixed to prepare FFPE samples. Resulting sections were used for IHC staining against specific antigens or subject to hematoxylin/eosin staining.

For chemoresistance studies, animals received subcutaneous implantation of tissue recombinants as described above and were given standard laboratory diets for 2 weeks to allow tumor uptake and growth initiation. Starting from the 3^rd^ week (tumors reaching 4-8 mm in diameter), MIT (0.2 mg/kg doses), DOX (doxorubicin, 1.0 mg/kg doses), therapeutic agent PDK4-IN (1.0 mg/kg doses, 200 μl/dose) or vehicle controls was administered by body injection (chemicals via intraperitoneal route, antibodies through tail vein), on the 1^st^ day of 3^rd^, 5^th^ and 7^th^ weeks, respectively. Upon completion of the 8-week therapeutic regimen, animals were sacrificed, with tumor volumes recorded and tissues processed for histological evaluation.

At the end of chemotherapy and/or targeting treatment, animals were anaesthetized and peripheral blood was gathered *via* cardiac puncture. Blood was transferred into a 1.5 ml Eppendorf tube and kept on ice for 45 min, followed by centrifugation at 9000 x g for 10 min at 4゜C. Clear supernatants containing serum were collected and transferred into a sterile 1.5 ml Eppendorf tube. All serum markers were measured using dry-slide technology on IDEXX VetTest 8008 chemistry analyzer (IDEXX). About 50 μl of the serum sample was loaded on the VetTest pipette tip followed by securely fitting it on the pipettor and manufacturer’s instructions were followed for further examination.

All animal experiments were performed in compliance with NIH Guide for the Care and Use of Laboratory Animals (National Academies Press, 2011) and the ARRIVE guidelines, and were approved by the Institutional Animal Care and Use Committee (IACUC) of Shanghai Institute of Nutrition and Health, Chinese Academy of Sciences.

#### Statistics

All *in vitro* experiments were performed in triplicates, while animal studies were conducted with at least 10 mice *per* group. Data are presented as mean ± SD except where otherwise indicated. GraphPad Prism 8.4.3 was used to collect and analyze data, with statistical significance determined according to individual settings. Cox proportional hazards regression model and multivariate Cox proportional hazards model analysis were performed with statistical software SPSS. Statistical significance was determined by unpaired two-tailed Student’s *t* test, one- or two-way ANOVA, Pearson’s correlation coefficients test, Kruskal-Wallis, log-rank test, Wilcoxon-Mann-Whitney test or Fisher’s exact test. For all statistical tests, a *P* value < 0.05 was considered significant.

To determine sample size, we began by setting the values of type I error (α) and power (1-β) to be statistically adequate: 0.05 and 0.80, respectively ^73^. We then determine *n* on the basis of the smallest effect we wish to measure. If the required sample size is too large, we chose to reassess the objectives or to more tightly control the experimental conditions to reduce the variance. We did not exclude samples or animals, nor statistical methods used to predetermine sample sizes, which, however, were similar to those generally employed in relevant fields.

## Data availability

Data that support the plots within this paper and other findings of this study are available from the corresponding author upon reasonable request. The RNA-seq data generated in the present study have been deposited in the Gene Expression Omnibus database under accession codes GSE198110.

## Acknowledgements

We are grateful to the members of Sun laboratory for reagents, comments and other contributions to this project. The work was supported by grants from National Key Research and Development Program of China (2020YFC2002800 and 2016YFC1302400 to Y.S.; 2019YFA0904800 to Y.Z.), National Natural Science Foundation of China (NSFC) (81472709, 31671425, 31871380, 82130045 to Y.S.; 32150030, 32030065, 32121005, 92049304 to Y.Z.; 81370730, 81571512 to Q.F.; 22007006 to G.Z.); the Strategic Priority Research Program of Chinese Academy of Sciences (XDB39010500) to Y.S.; Shanghai Municipal Science and Technology Commission Excellent Academic Leader Program (20XD1404300) to Y.S.; Anti-Ageing Collaborative Program of SIBS and BY-HEALTH (C01201911260006, C01202112160005) to Y.S.; the University and Locality Collaborative Development Program of Yantai (2019XDRHXMRC08 and 2020XDRHXMXK02 to Y.S.; 2021XDHZ082 to Q.F.); Natural Science Foundation of Shandong Province Collaborative Fund (ZR2021LSW021) to Y.S.; and the U.S. DoD PCRP (Idea Development Award PC111703) to Y.S.; Research Unit of New Techniques for Live-cell Metabolic Imaging (Chinese Academy of Medical Sciences, 2019-I2M-5-013 to Y.Z.), Innovative research team of high-level local universities in Shanghai (Y.Z.); Yantai Double Hundred Program to Q.F.; Taishan Scholars Construction Engineering (tsqn201909144), Special Project of Central Government for Local Science and Technology Development of Shandong Province (YDZX20203700001291) to G.Z.

## Author contributions

Y.S. conceived this study, designed the experiments and orchestrated the project. X.D. performed most of the *in vitro* assays, part of the *in vivo* experiments and wrote part of the manuscript. Q.L and D.F. acquired and analyzed clinical samples from prostate and breast cancer patients, respectively, and managed subject information. Y.Z. and S.L. helped with metabolic profiling and GC-MS analysis of cell metabolites with SoNar- and FiLa-based genetically encoded fluorescence sensors. X.C., Q.X. and C.W. performed some cell culture and drug treatment assays. G.Z. provided constructive advices. Q.F. performed partial preclinical studies. Y.Z. and J.C. provided conceptual inputs and/or supervised a specific subset of experiments. Y.Z. organized the generation of SoNar-and FiLa-associated graphic illustrations. Y.S. performed data analysis, graphic presentation and finalized the manuscript. All authors critically read and commented on the final manuscript.

## Inclusion & Ethics statement

All authors have agreed to all manuscript contents, the author list and its order and the author contribution statements. Any changes to the author list after submission will be subject to approval by all authors.

## Additional information

**Supplementary information** The online version contains supplementary material available at specific link.

**Correspondence** and requests for materials should be addressed to Yu Sun.

**Reprints and permissions information** is available online.

