## Supplemental figures for "Senescent cells develop PDK4-dependent hypercatabolism and form an acidic microenvironment to drive cancer resistance"

### Supplementary Fig. 1

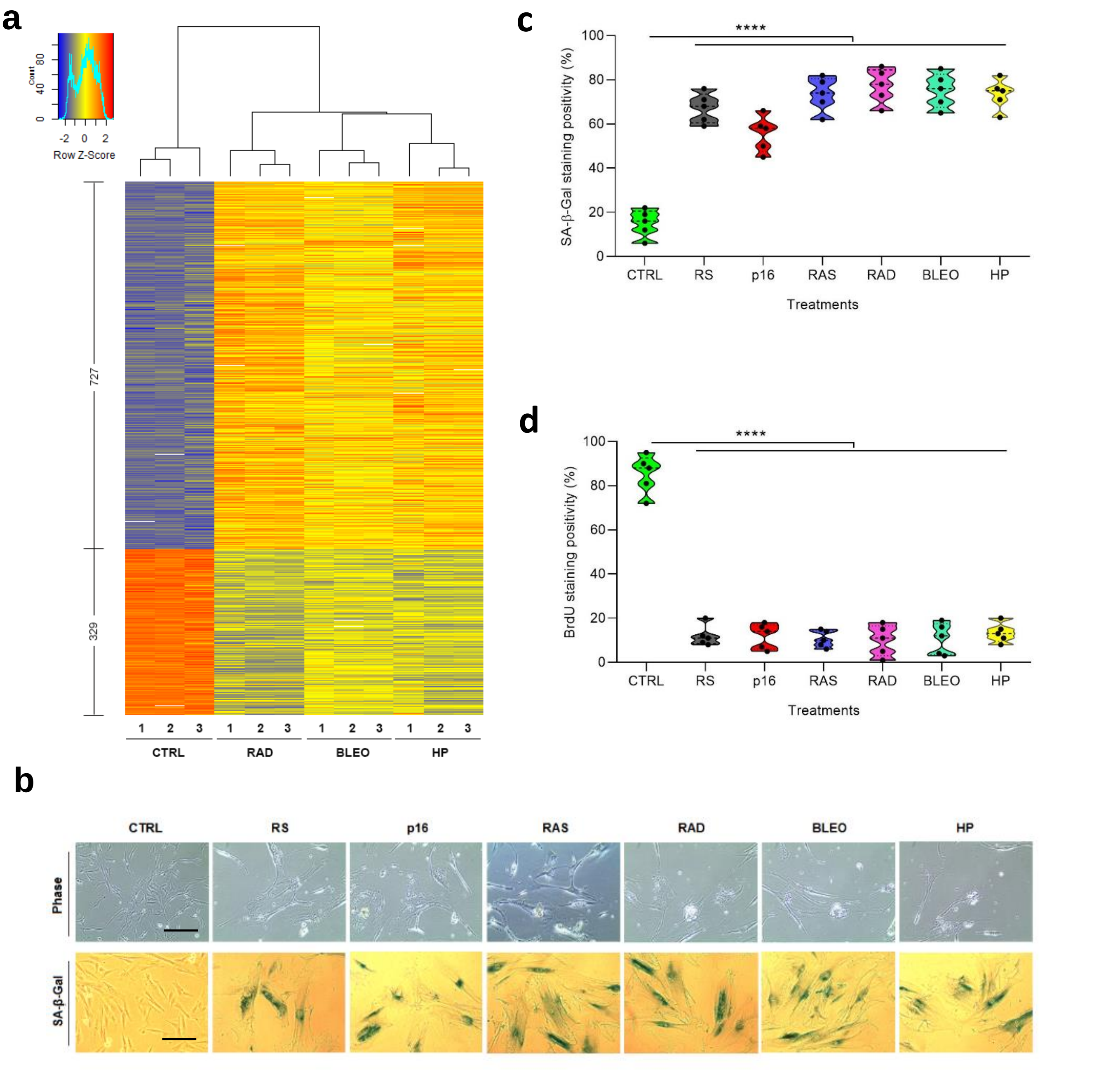

### Supplementary Fig. 2

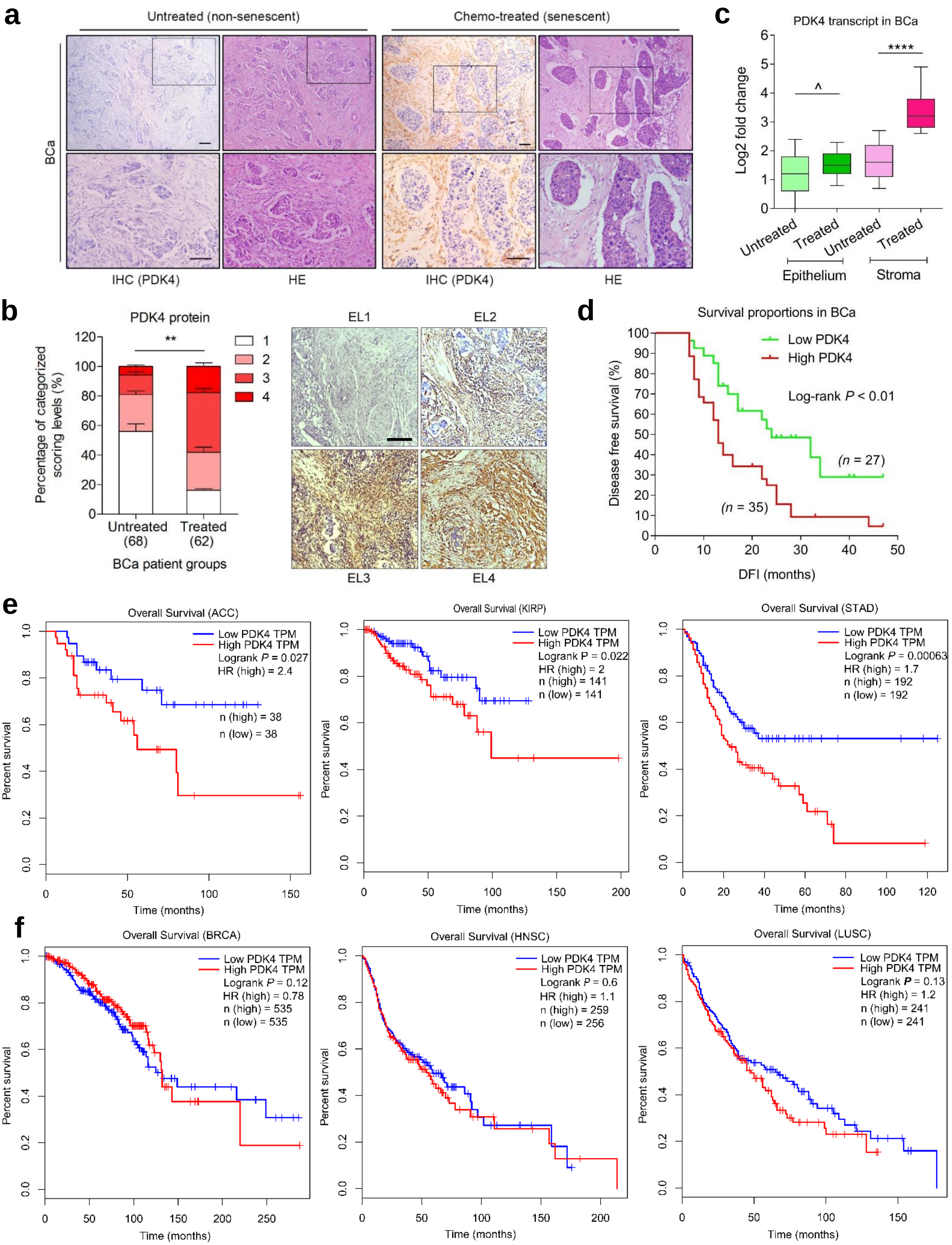

#### Supplementary Fig. 3

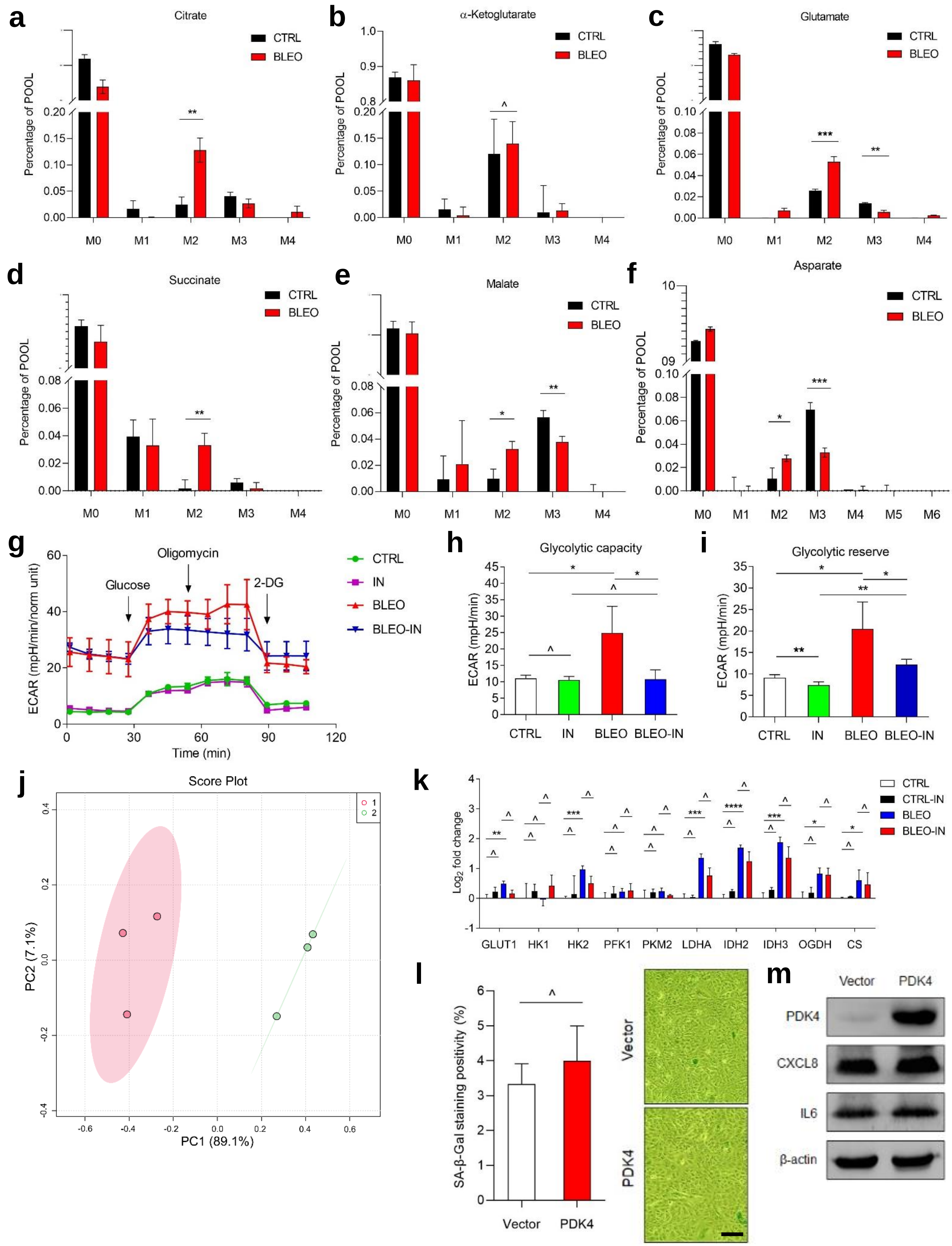

### Supplementary Fig. 4

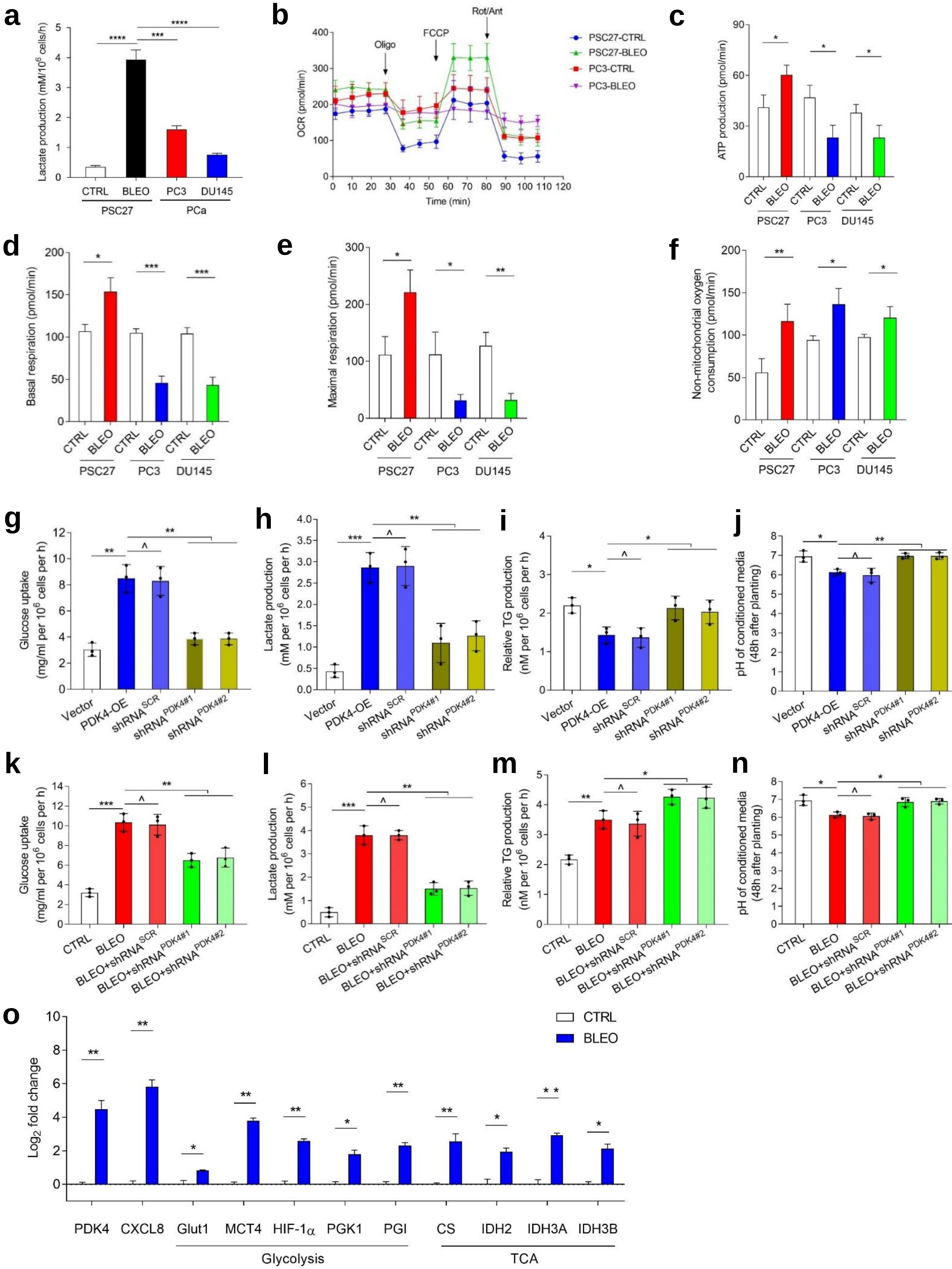

### Supplementary Fig. 5

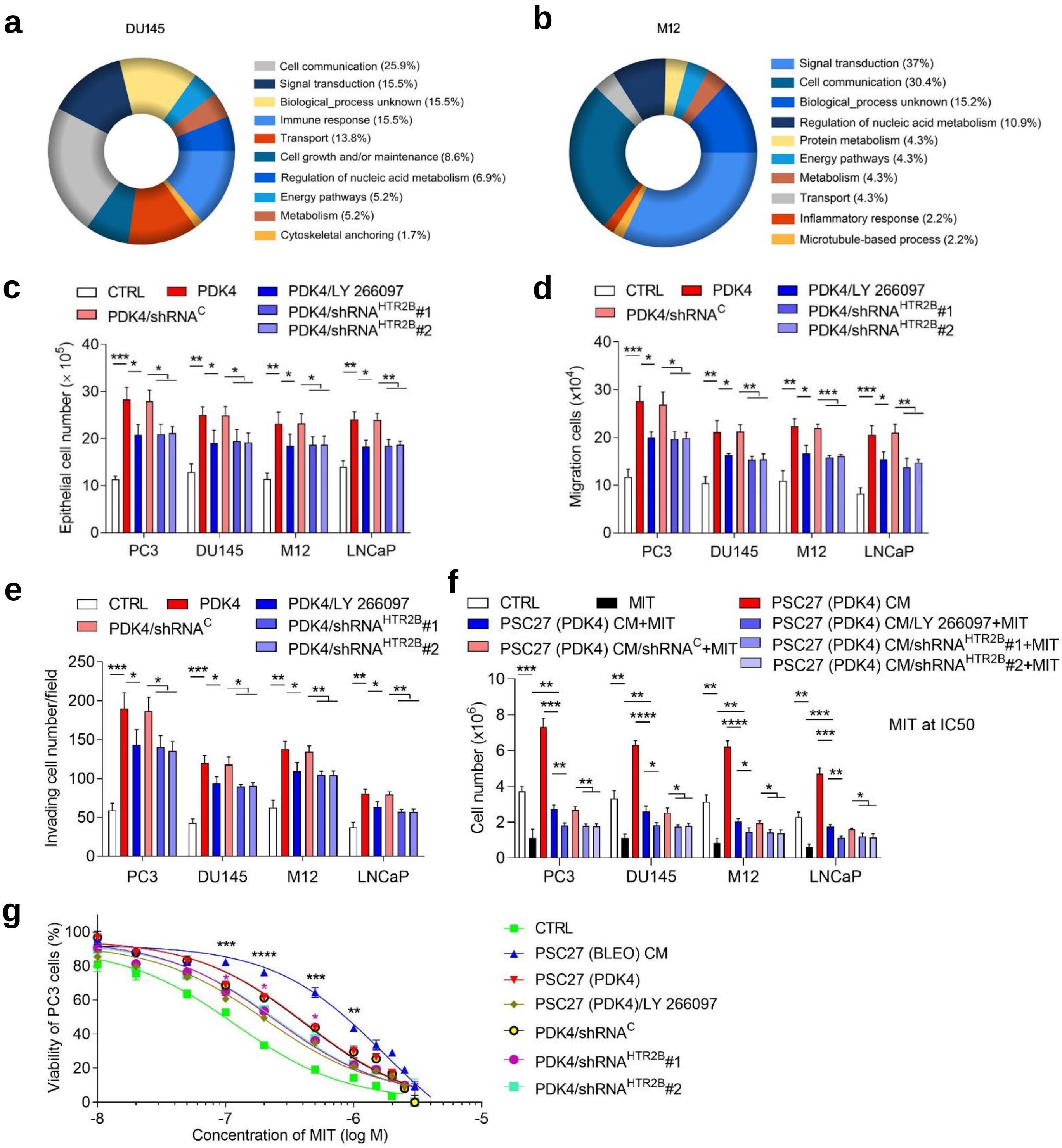

### Supplementary Fig. 6

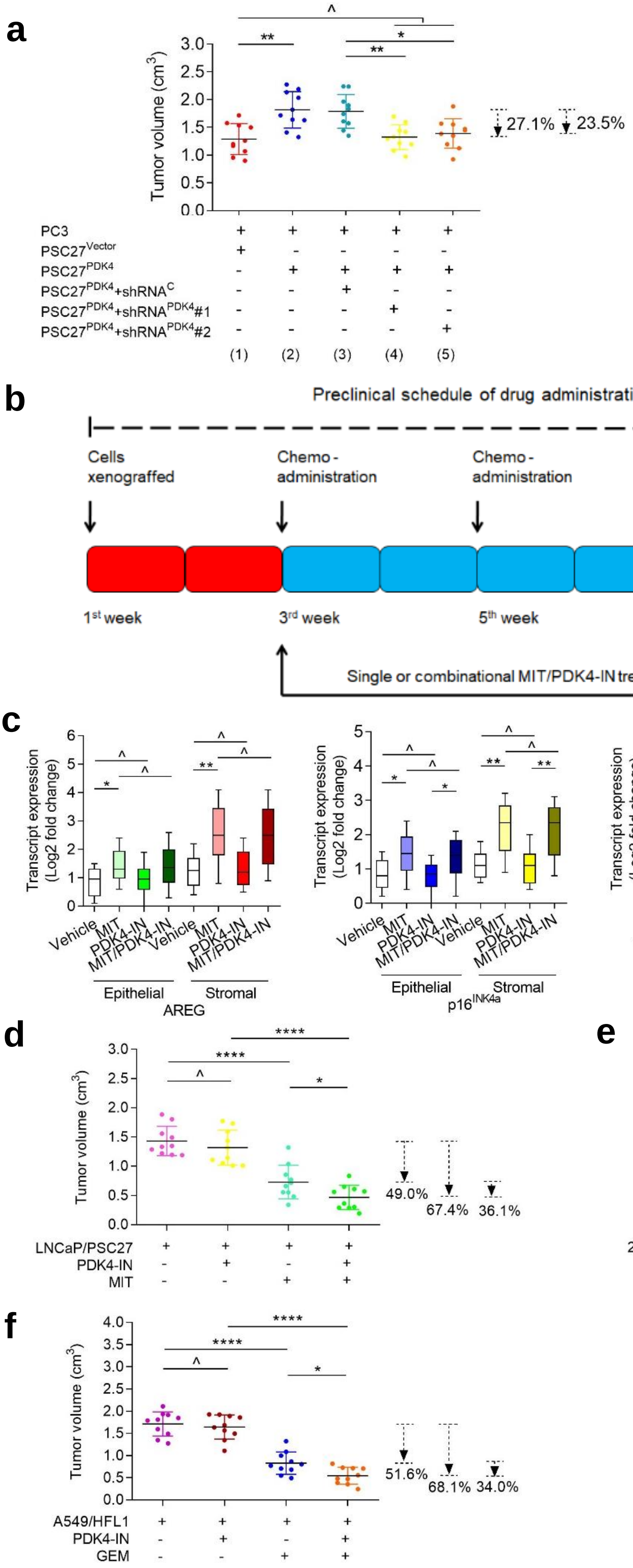

#### Supplementary Fig. 7

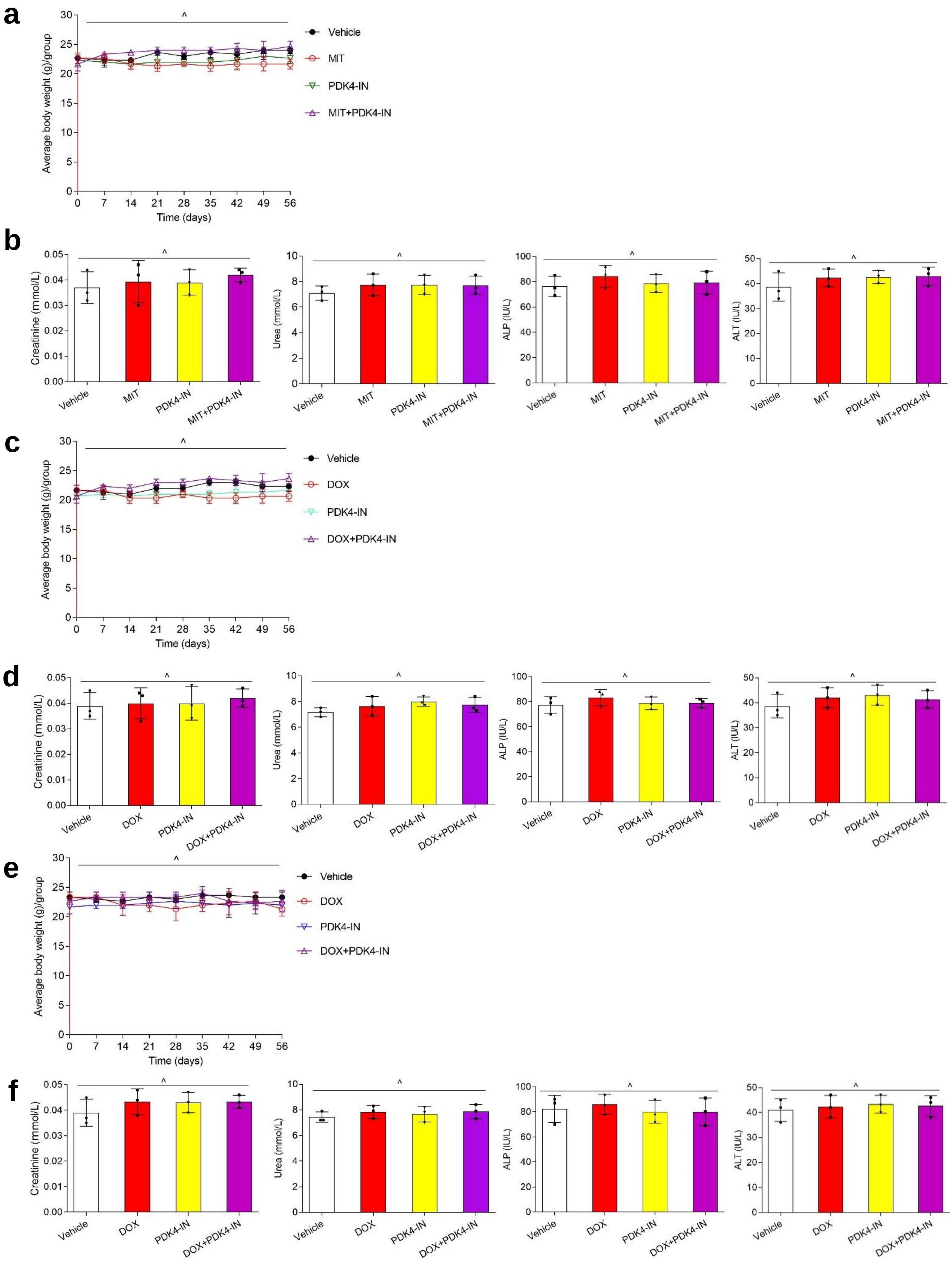
